# ER-to-Golgi transport machinery promotes the excessive cargo-triggered unfolded protein response

**DOI:** 10.64898/2025.12.15.694336

**Authors:** Liying Guan, Tong Zhang, Zhigao Zhan, Yingchun Wang, Xun Huang, Mei Ding

## Abstract

Disruptions to ER homeostasis trigger the unfolded protein response (UPR) to restore proteostatic balance. While defects in the secretory machinery are known to induce ER stress, it remains unclear whether specific trafficking components directly modulate UPR signaling. Here, we demonstrate that neuronal overexpression of the gap junction protein UNC-9 activates the IRE1-XBP1 arm of the UPR in *C. elegans*. Genetic deletion of *ERGIC2* or *ERGIC3*—genes encoding COPII-associated proteins required for UNC-9 transport—suppresses this UPR activation, revealing an unexpected role for these factors beyond cargo trafficking. Mechanistically, ERGIC2 and ERGIC3 interact with the ER chaperone BiP, facilitating its release from IRE1 to enable UPR and alleviate cargo aggregation. Our findings redefine the UPR as a process dynamically regulated by early secretory components and provide novel insights into how cells integrate trafficking demand with stress adaptation, with implications for ER stress-associated diseases such as neurodegeneration.

## Introduction

The endoplasmic reticulum (ER) is the central hub for protein folding, quality control, and secretion, processing nearly one-third of the proteome. Properly folded proteins are packaged into COPII vesicles for transport to the Golgi, while misfolded proteins are targeted for ER-associated degradation (ERAD) (Araki and Nagata, 2011; Ellgaard and Helenius, 2003; Plate and Wiseman, 2017). Disruptions in secretory trafficking—due to genetic mutations, environmental stress, or excessive protein synthesis—can lead to protein aggregation and ER dysfunction, triggering the unfolded protein response (UPR) to restore homeostasis (Mercado et al., 2013; Sicari et al., 2019; Tang, 2021; Zhang et al., 2021). While deficiencies in secretory machinery are known to induce ER stress and UPR activation, however, it is largely unknown whether the secretory machinery itself directly modulates UPR signaling.

In metazoans, the UPR is mediated by three ER transmembrane sensors: IRE1(inositol requiring enzyme 1), PERK (double-stranded RNA-activated protein kinase [PKR]-like ER kinase), and ATF6 (activating transcription factor 6) (Hetz and Saxena, 2017; Walter and Ron, 2011). Among these, IRE1 is the most evolutionarily conserved and, upon ER stress, oligomerizes to splice *XBP1* mRNA, producing the active transcription factor XBP1s (Calfon et al., 2002; Cox and Walter, 1996; Yoshida et al., 2001). This drives the expression of chaperones and ERAD components, enhancing protein-folding capacity and clearance of misfolded proteins (Ron and Walter, 2007). While extensive research has elucidated UPR activation mechanisms and their links to disease, the potential regulatory role of secretory cargo burden, the flux of proteins transiting through the ER-Golgi pathway, remains poorly understood.

Here, we demonstrate that neuronal overexpression of four-transmembrane gap junction protein UNC-9 triggers the IRE1-XBP1-medaited UPR, but surprisingly, loss of *ERGIC2* or *ERGIC3*—genes encoding COPII-associated proteins required for UNC-9 transport (Guan et al., 2022; Orci et al., 2003; Shibuya et al., 2015; Welsh et al., 2006) —inhibits this UPR activation. Mechanistically, ERGIC2 and ERGIC3 interact with the ER chaperone BiP, promoting its dissociation from IRE1 and enabling UPR activation. This finding challenges the conventional view of ERGIC2 and ERGIC3 as mere cargo transporters, instead implicating them as active regulators of UPR signaling. Together, our work provided novel insights into how cells integrate protein trafficking demands with stress resilience, with broad implications for ER stress-related diseases, including neurodegeneration and metabolic disorders.

## Results

### *ergi-2* and *ergi-3* are required for the ER stress response induced by UNC-9 overexpression

In *C. elegans*, the D-type motor neurons (DD/VD) have their cell bodies located on the ventral nerve cord. They send processes across the worm’s body to the dorsal cord; from there, the processes branch out and extend ventrally and dorsally, respectively (Fig. 1A) (White et al., 1986). *hsp-4* encodes the worm homolog of the ER chaperone BiP, which is transcriptionally upregulated during ER stress (Calfon et al., 2002). Under basal conditions, an *hsp-4* promoter-driven mCherry reporter (P*hsp-*4::mCherry) exhibits weak expression across intestinal and epidermal cells (Fig. 1B) (Burga et al., 2011; Guan et al., 2020). When we used the P*unc-25* promoter to drive expression of the four-transmembrane gap junction protein UNC-9 specifically in DD/VD neurons (Jin et al., 1999), the P*hsp-*4::mCherry signal increased dramatically and specifically in these neurons (indicated by asterisks and arrows in Fig. 1B) (Guan et al., 2020). This suggests that UNC-9 overexpression induces an ER stress response in DD/VD neurons.

**Figure 1:**
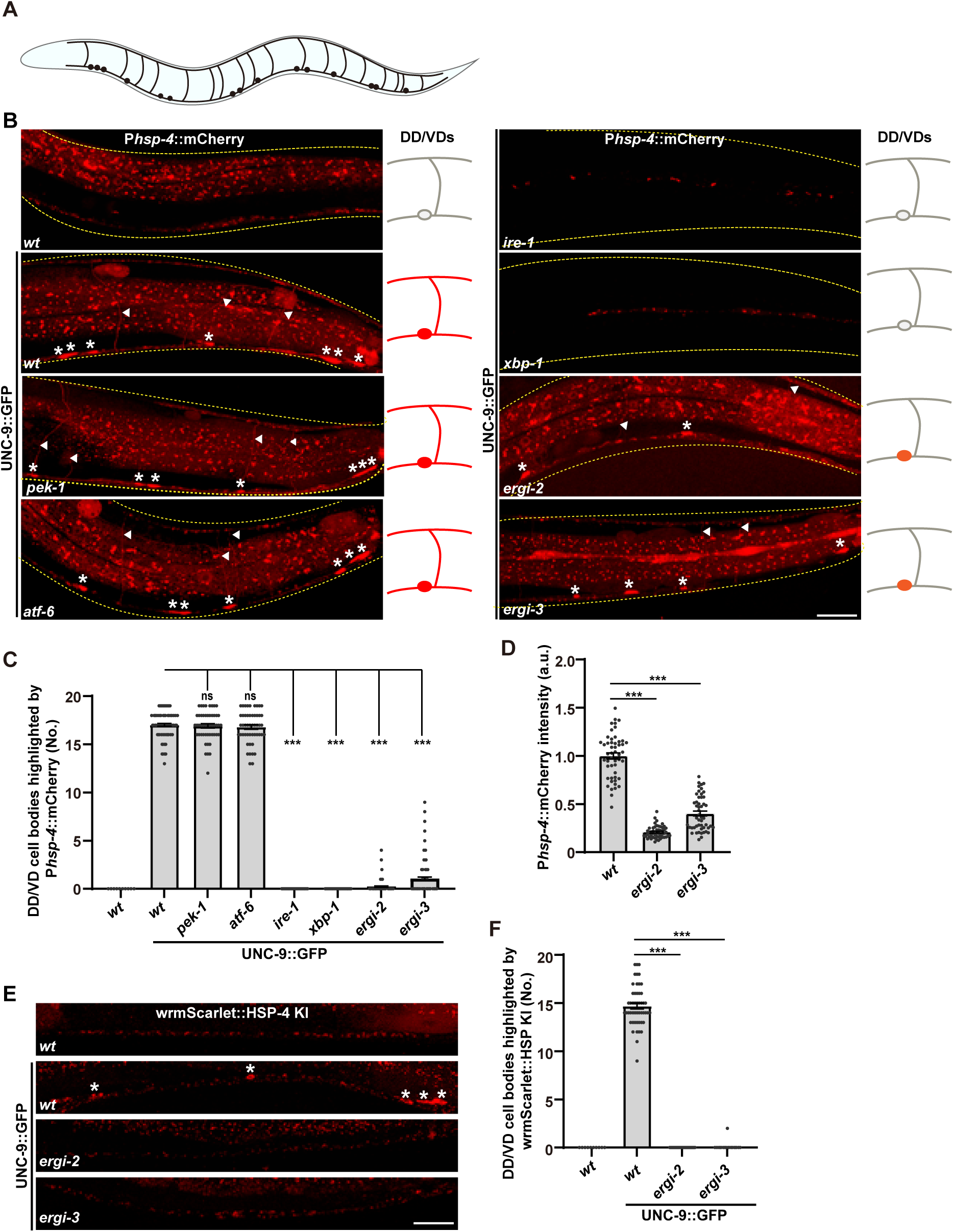
ERGI-2 or ERGI*-*3 depletion inhibits the UPR. (A) Schematic of DD/VD neurons in *C. elegans.* (B) UNC-9::GFP (green) induces P*hsp-4*::mCherry (red) expression, which is reduced in *ire-1, xbp-1, ergi-2* and *ergi-3* mutants, but not in *pek-1* and *atf-6* mutants. Yellow dashed lines outline the worms. White triangles indicate commissures; asterisks mark cell bodies. Scale bar: 25μm. (C) Quantification of mCherry-positive DD/VD cell bodies. (D) Quantification of mCherry fluorescence intensity in cell bodies. (E) UNC-9::GFP (green) induces wrmScarlet::HSP-4 KI (red) expression, which is inhibited in *ergi-2* and *ergi-3* mutants. Asterisks mark cell bodies. Scale bar: 25μm. (E) Quantification of wrmScarlet-positive DD/VD cell bodies. Data are mean ± SEM. One-way ANOVA with Tukey’s test was used. ***P <0.001; ns, not significant.

The ER stress response can be mediated by the IRE1, PERK or ATF6 pathways (Walter and Ron, 2011). Introducing *pek-1* or *arf-6* mutation into UNC-9 overexpression worms did not affect *hsp-4* induction (Fig. 1B and C). In contrast, mutations in *ire-1* or *xbp-1* greatly reduced *hsp-4* induction (Fig. 1B and C), indicating that the IRE1-XBP1 branch is specifically required for the UNC-9 overexpression-triggered UPR. Noticeably, the basal expression of *hsp-4* was reduced by *ire-1* or *xbp-1* mutation, suggesting that basal *hsp-4* expression also requires the IRE1-XBP1 branch.

To understand how the IRE1-XBP1 branch is regulated *in vivo*, we searched for mutants displaying reduced *hsp-4* induction similar to *ire-1* or *xbp-1* and identified *ergi-2* and *ergi-3*. In *ergi-2* or *ergi-3* mutant animals, the UNC-9-induced P*hsp-4*::mCherry upregulation was significantly reduced (Fig. 1B-D). Both the fraction of mCherry-positive DD/VD neurons and their fluorescence intensity were markedly reduced compared to wild type (Fig. 1B-D). This reduction was not due to neuronal loss, as the number of DD/VD neurons remained unchanged (Sup Fig. 1A and B).

We further determined whether endogenous *hsp-4* gene expression is similarly regulated by *ergi-2* and *ergi-3*. For this, a wrmScarlet::HSP-4 knock-in strain was generated. Overexpression of UNC-9::GFP robustly induced the wrmScarlet::HSP-4 signal in DD/VD neurons (Fig. 1E and F), confirming that endogenous *hsp-4* expression is upregulated by excessive UNC-9. Noticeably, in *ergi-2* or *ergi-3* mutants, this induction was largely suppressed (Fig. 1E and F). Thus, these results demonstrate that *ergi-2* and *ergi-3* are required for the ER stress response triggered by UNC-9 overexpression in DD/VD neurons.

### *ergi-2* and *ergi-3* function upstream of *ire-1* in the UPR pathway

UNC-9 overexpression in DD/VD neurons selectively activates the IRE1-XBP1 branch of the UPR. In addition to *hsp-4* induction, this pathway also activates autophagy, providing a second readout for UPR activation (Guan et al., 2020). To determine whether *ergi-2* and *ergi-3* mediate their effects through the IRE1-XBP1 axis, we examined autophagy induction using a DD/VD-specific mCherry::LGG-1 reporter (*C. elegans* Atg8). Under basal conditions, mCherry::LGG-1 exhibited diffuse cytoplasmic localization (Fig. 2A and B) (Guan et al., 2020). Upon UNC-9::GFP overexpression, a dramatic increase in mCherry::LGG-1 puncta appeared, confirming autophagy activation (Fig. 2A and B). In *ergi-2* and *ergi-3* mutants, autophagy induction was strongly suppressed, with mCherry::LGG-1 puncta formation reduced to near-baseline levels (Fig. 2A and B). Therefore, *ergi-2* and *ergi-3* are also required for autophagy activation induced by UNC-9::GFP overexpression.

**Figure 2:**
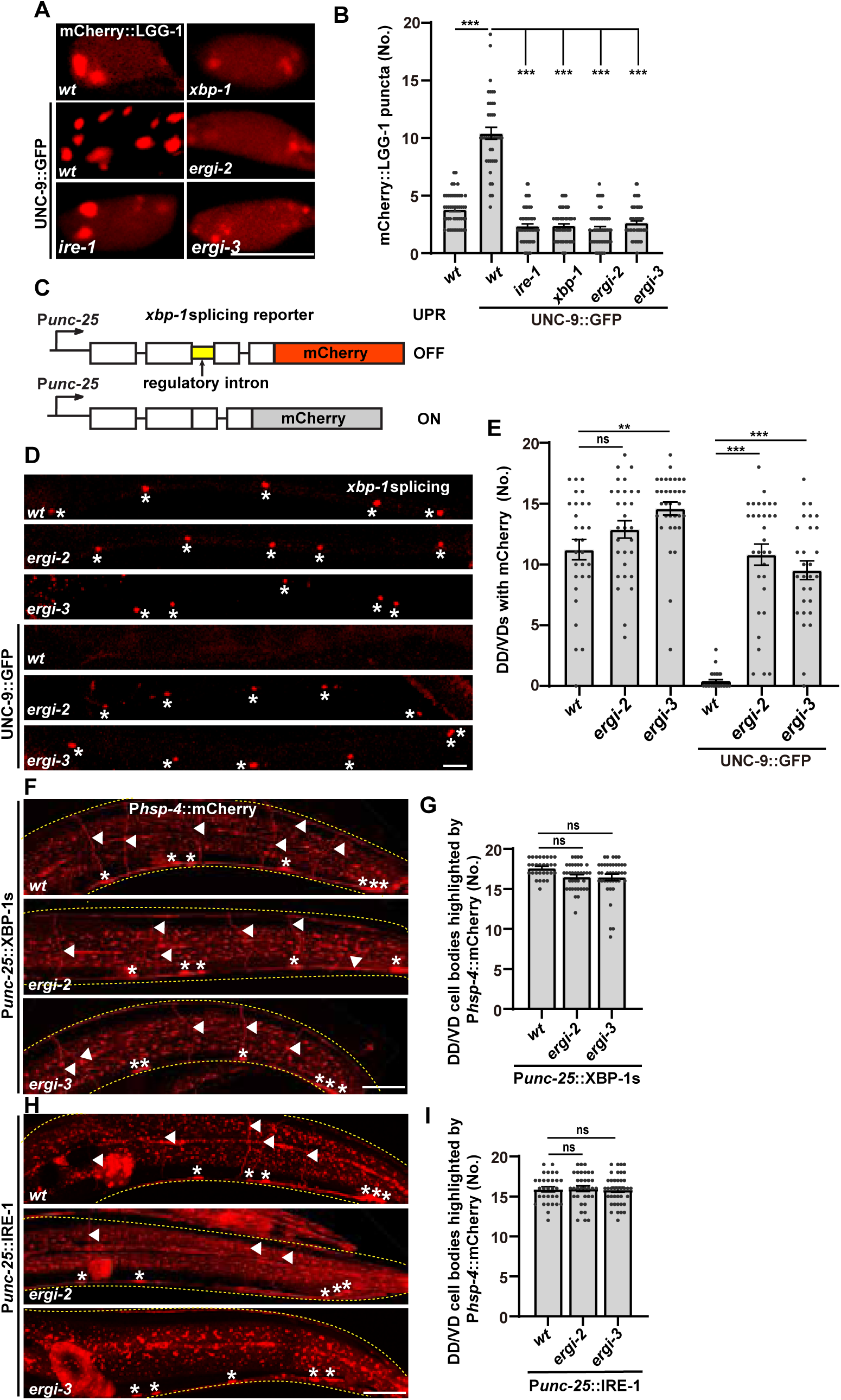
*ergi-2* and *ergi-*3 function upstream of *ire-1*. (A) UNC-9::GFP induces mCherry::LGG-1 (red) puncta formation, which is inhibited in *ire-1, xbp-1, ergi-2* and *ergi-3* mutants. Scale bar: 5μm. (B) Quantification of mCherry::LGG-1 puncta number in cell bodies. (C) Schematic of the *xbp-1* splicing reporter. (D) UNC-9::GFP induces *xbp-1* mRNA splicing which is inhibited in *ergi-2* and *ergi-3* mutants. Asterisks mark cell bodies. Scale bar: 25μm. (E) Quantification of mCherry-positive DD/VD cell bodies from (D). (F) XBP-1s overexpression induces P*hsp-4*::mCherry (red), which remains in *ergi-2* and *ergi-3* mutants. (G) Quantification of mCherry-positive DD/VD cell bodies from (F). (H) IRE-1 overexpression induces P*hsp-4*::mCherry (red), which remains in *ergi-2* and *ergi-3* mutants. (I) Quantification of mCherry-positive DD/VD cell bodies from (H). In (F) and (H), white triangles indicate commissures; asterisks mark cell bodies; scar bars: 25μm. Data are mean ± SEM. One-way ANOVA with Tukey’s test was used. **P <0.01; ***P <0.001; ns, not significant.

A key event in the IRE1-XBP1 pathway is IRE1-dependent splicing of *xbp-1* mRNA. To monitor *xbp-1* mRNA splicing in DD/VD neurons, we developed an *in vivo* reporter. (Guan et al., 2020; Shim et al., 2004). This reporter consists of mCherry fused to the un-spliced (inactive) form of *xbp-1*, driven by the DD/VD specific promoter P*unc-25* (Fig. 2C). In the absence of ER stress, the unprocessed transcript produces mCherry expression (“mCherry on”). Upon UPR activation, IRE1-mediated splicing removes the intron, abolishing mCherry expression (“mCherry off”) (Fig. 2C). Consistent with our earlier findings, UNC-9::GFP overexpression induced robust splicing (“mCherry off”), reflecting pathway activation. In *ergi*-2 and *ergi-3* mutants, we found that the *xbp-1* splicing was severely impaired, as evidenced by persistent mCherry signal in DD/VD neurons (Fig. 2D and E). This result places *ergi-2* and *ergi-3* upstream of *xbp-1* splicing, implicating them in the initiation or propagation of the IRE1-dependent UPR signal. Noticeably, without UNC-9::GFP overexpression, the absence of *ergi-2* or *ergi-3* alone did not induce *xbp-1* splicing (Fig. 2D and E), suggesting that *ergi-2* and *ergi-3* are not involved in UPR activation in general.

Overexpression of either the spliced form of XBP-1 (XBP-1s) or IRE-1 in DD/VD neurons is sufficient to activate the UPR, indicated by P*hsp-*4::mCherry induction (Fig. 2F-I). Interestingly, this induction remained unaffected in *ergi-2* or *ergi-3* mutants (Fig. 2F-I), suggesting that *ergi-2* and *ergi-3* may act upstream of *ire-1* in the pathway leading to UPR activation.

### The specific role of *ergi-2* and *ergi-3* in UPR activation

*ergi-2* and *ergi-3* encode the worm homologies of yeast Erv41 and Erv46, and mammalian ERGIC2 and ERGIC3, respectively. Previous studies indicate that the Erv41/ERGIC2 and Erv46/ERGIC3 proteins are associated with COPII complex and play a regulatory role in ER-to-Golgi secretory pathway (Adolf et al., 2019; Barlowe, 2002; Guan et al., 2022; Keiser and Barlowe, 2020; Shibuya et al., 2015; Stefan Otte, 2001; Welsh et al., 2006). To understand how *ergi-2* and *ergi-3* function in UPR activation, we tested whether COPII components are involved in the UNC-9-induced stress response using a tissue-specific RNAi approach. Knocking down core COPII components (SEC-12 or SEC-13) in DD/VD neurons resulted in significant neuronal cell loss, consistent with the essential role of the COPII complex for viability (Sup Fig. 1A and B). In the remaining DD/VD neurons, the P*hsp-4::*mCherry signal intensity was similar to controls (Sup Fig. 1C and D). This suggests that while the COPII pathway is essential for DD/VD neuron viability, it is not required for UNC-9 overexpression-induced UPR activation.

ERGIC32 shares structural similarity with ERGIC2 and ERGIC3 and has been shown to bind to ERGIC3 and regulates its subcellular localization (Breuza et al., 2004). However, in *ergi-1* (the *C. elegans* homolog of ERGIC32) mutant animals, UNC-9-triggered UPR activation was indistinguishable from wild type (Sup Fig. 1E and F). ERGIC53, another component of the ER-Golgi intermediate compartment critical for cargo transport (Hauri et al., 2000; Schindler et al., 1993), was also dispensable for UNC-9-induced stress response, as *ERGIC53/ile-1* mutants exhibited wild-type levels of P*hsp-4::*mCherry induction (Sup Fig. 1H and I).

Together, these findings indicate that *ergi-2* and *ergi-3* play a specific role in mediating excessive UNC-9-induced ER stress response, independent of general ER-to-Gologi trafficking machinery.

### *ergi-2* and *ergi-3* are specifically required for excessive UNC-9-induced UPR

Treatment with tunicamycin (Tm), an N-linked glycosylation inhibitor that disrupts protein folding, robustly activated the IRE1-XBP1 branch of the UPR in *C. elegans* (Bar-Ziv et al., 2020; Calfon et al., 2002), This was evidenced by a strong induction of the ER stress reporter P*hsp-4*::mCherry in the intestine (Fig. 3A and B).

**Figure 3:**
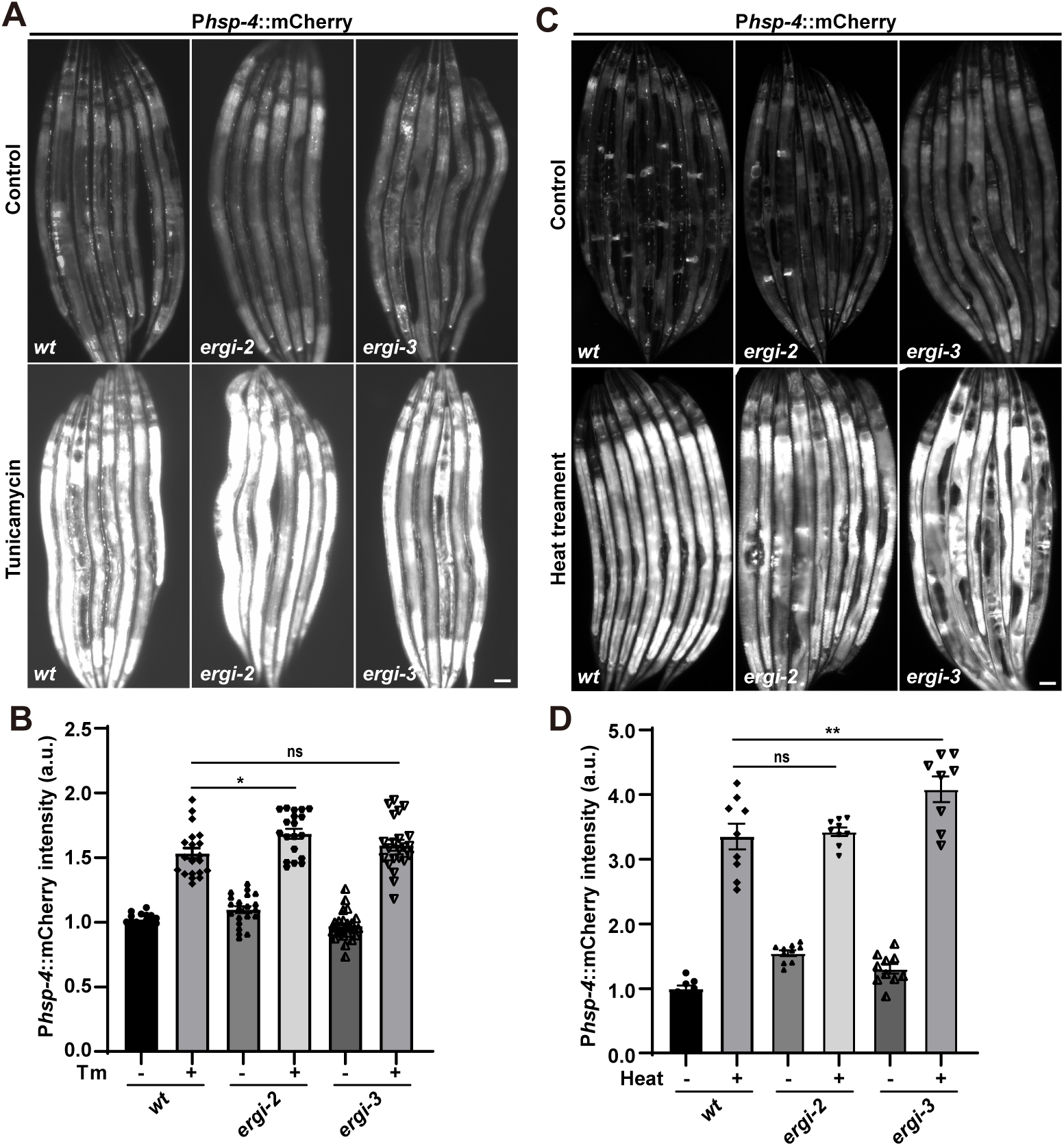
*ergi-2* and *ergi-3* do not affect the UPR triggered by tunicamycin or heat treatment. (A) Tunicamycin treatment induces P*hsp-4*::mCherry expression, which is unaffected in *ergi-2* and *ergi-3* mutants. (B) Quantification of P*hsp-4*::mCherry intensity in the intestine region for (A). (C) Heat treatment induces P*hsp-4*::mCherry expression, which is unaffected in *ergi-2* and *ergi-3* mutants. (D) Quantification of P*hsp-4*::mCherry intensity in the intestine region for (C). Scale bars: 25μm. Data are mean ± SEM. One-way ANOVA with Tukey’s test was used. **P <0.01; ***P <0.001; ns, not significant.

Surprisingly, loss-of-function mutations in *ergi-2* or *ergi-3* did not attenuate this Tm-induced P*hsp-4*::mCherry upregulation (Fig. 3A and B). Heat shock, another well-characterized stressor, also elicited intestinal P*hsp-4*::mCherry expression (Fig. 3C and D) (Al-Amin et al., 2016; Zhou et al., 2024), and this response remained intact in *ergi-2* or *ergi-3* mutants (Fig. 3C and D). Therefore, both *ergi-2* and *ergi-3* are dispensable for the UPR triggered by global ER protein misfolding.

Since Tunicamycin- or heat shock-induced UPR does not require *ergi-2* or *ergi-3*, we investigated whether these genes play a specific role in neuronal UPR activation triggered by misfolded or overexpressed secretory proteins. To this end, we examined two distinct secretory proteins: UNC-6/Netrin, an evolutionarily conserved extracellular guidance cue for neuronal migration and axon pathfinding (Asakura et al., 2015), and VIT-2, a yolk core protein involved in lipid transport (Zhai and Dong, 2025). Overexpression of UNC-6::GFP or VIT-2::GFP in DD/VD neurons led to significant upregulation of *Phsp-4*::mCherry specifically in these neurons, confirming UPR activation due to secretory protein overload (Fig. 4A-D). However, mutations in *ergi-2* or *ergi-3* did not suppress this induction (Fig. 4A–D), indicating that these genes are not broadly required for UPR activation by protein overexpression in neurons.

**Figure 4:**
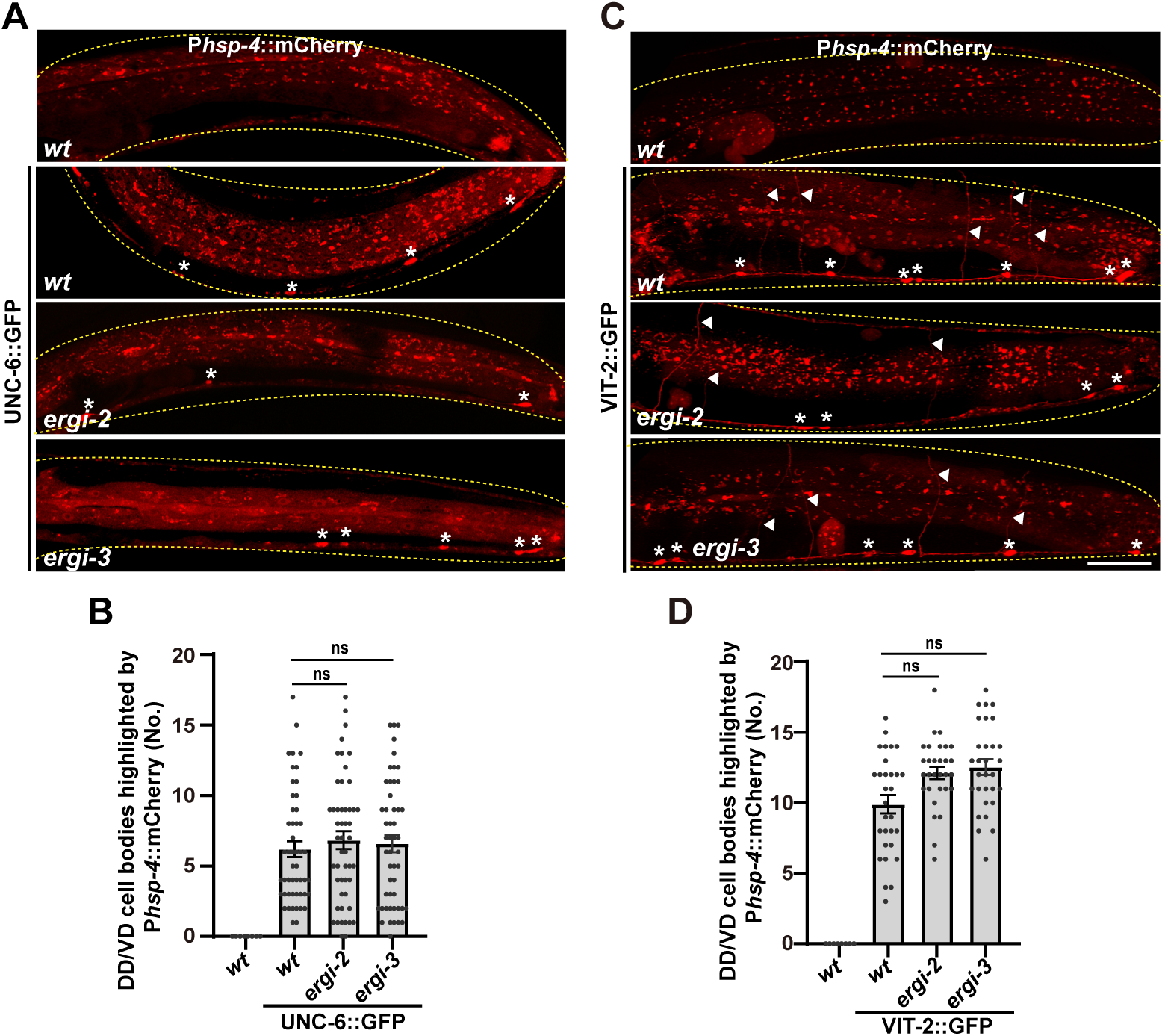
*ergi-2* and *ergi-3* do not affect the UPR triggered by neuronal overexpression of UNC-6::GFP or VIT-2::GFP. (A) UNC-6::GFP expression in DD/VD neurons induces P*hsp-4*::mCherry, which is unaffected in *ergi-2* and *ergi-3* mutants. (B) Quantification of mCherry-positive DD/VD cell bodies from (A). (C) VIT-2::GFP expression in DD/VD neurons induces P*hsp-4*::mCherry, which is unaffected in *ergi-2* and *ergi-3* mutants. (D) Quantification of mCherry-positive DD/VD cell bodies from (C). Yellow dashed lines outline the worms. White triangles indicate commissures; asterisks mark cell bodies. Scale bars: 25μm. Data are mean ± SEM. One-way ANOVA with Tukey’s test was used. ns, not significant.

In contrast, the UNC-9 overexpression-triggered UPR was strongly dependent on *ergi-2* and *ergi-3*. This selectivity implies that ERGI-2 and ERGI-3 are not general UPR mediators but instead participate in a substrate-specific surveillance mechanism for certain misfolded or overloaded proteins.

### Excessive UNC-9 accumulates in the ER in e*rgi-2* and *ergi-3* mutants

To understand the role of ERGI-2 and ERGI-3 in the UPR specifically triggered by excessive UNC-9, we examined UNC-9 localization in DD/VD neurons. In wild type animals, ectopically expressed UNC-9::GFP forms puncta along neurites and within cell bodies (Fig. 5A) (Guan et al., 2020). The *ire-1* and *xbp-1* genes are essential for the UNC-9 overexpression-induced UPR. In *ire-1* or *xbp-1* mutants, UNC-9::GFP was diffusely distributed in DD/VD cell bodies, and the punctate UNC-9::GFP signal on neurites was greatly reduced (Fig. 5A). Double-labeling experiments confirmed that this diffuse UNC-9::GFP is retained in the ER (Fig. 5B) (Guan et al., 2020), consistent with the notion that UPR deficiency leads to defective folding and ER retention of overexpressed UNC-9.

**Figure 5:**
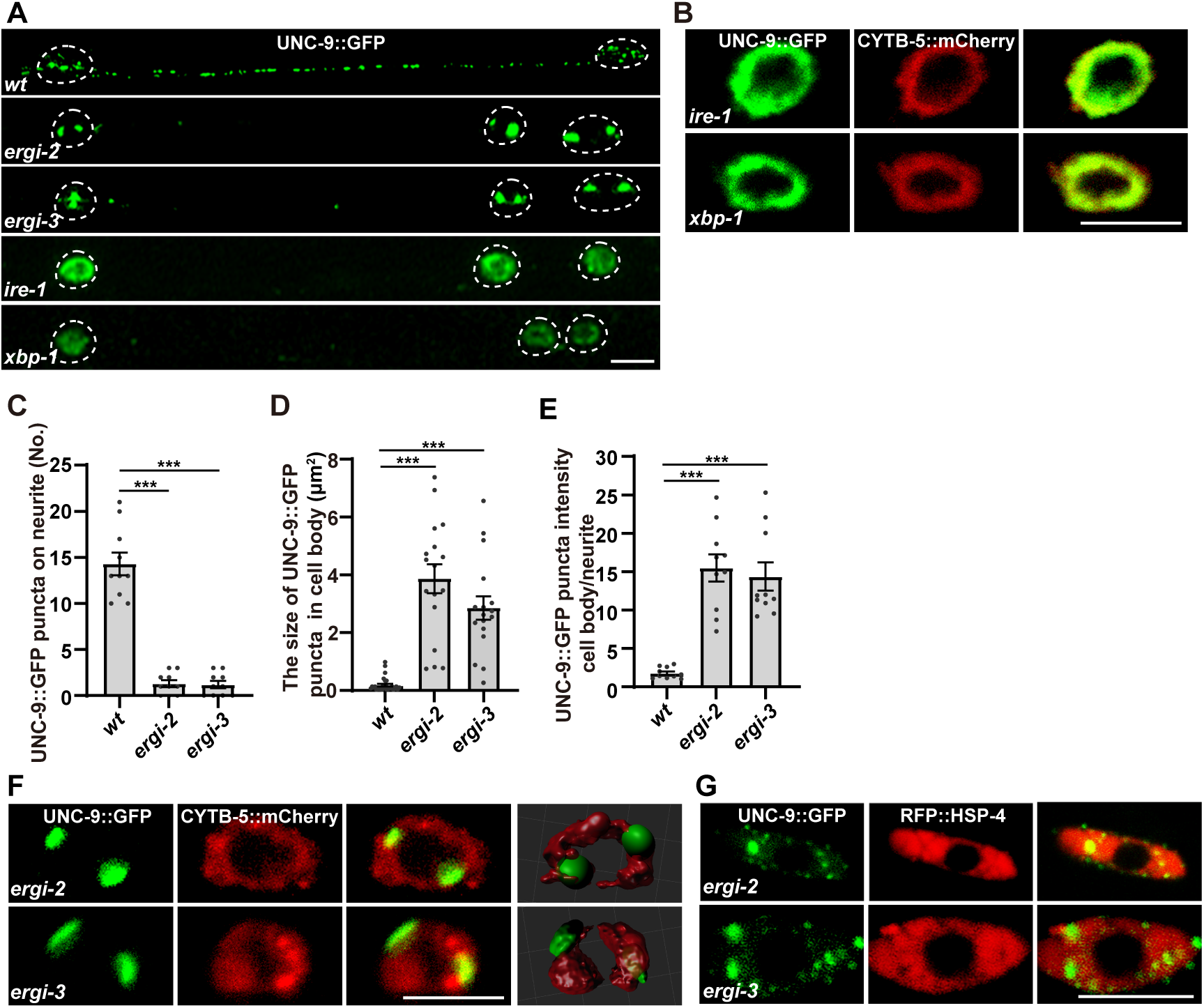
Large UNC-9::GFP aggregates accumulate in the ER in *ergi-*2 and *ergi-3* mutants. (A) The distribution of UNC-9::GFP (green) in DD/VD neuronal cell bodies and neurites in the indicated genotypes. (B) UNC-9::GFP (green) co-localizes with the ER markers CYTB-5::mCherry (red) in *ire-1* and *xbp-1* mutants. (C) Quantification of the number of UNC-9::GFP on neurites. (D) Quantification of the size of UNC-9::GFP puncta in DD/VD cell bodies. (E) Quantification of the fluorescence intensity of UNC-9::GFP in cell bodies compared to neurites. (F) Large UNC-9::GFP puncta are localized to the ER (marked with CYTB-5::mCherry, red) in *ergi-2* and *ergi-3* mutants. (G) UNC-9::GFP puncta co-localize with RFP::HSP-4 (red) in *ergi-2* and *ergi-3* mutants. Scale bar: 5μm. Data are mean ± SEM. One-way ANOVA with Tukey’s test was used. **P <0.01; ***P <0.001; ns, not significant.

In contrast, *ergi-2* and *ergi-3* mutations resulted in large UNC-9::GFP puncta accumulated in the DD/VD cell body region (Fig. 5A). Compared to wild type, the size of these puncta was considerably increased, while their number was significantly reduced (Fig. 5A and D). Concurrently, UNC-9::GFP signal on DD/VD neurites was greatly diminished (Fig. 5C and E).

We next sought to identify the subcellular compartment containing these enlarged UNC-9::GFP aggregates in *ergi-2* and *ergi-3* mutants. Our previous studies indicated that *ergi-2* or *ergi-3* are required for ER-to-Golgi transport of gap junction proteins (Guan et al., 2022). Double-labeling in DD/VD neurons with the ER marker CYTB-5::mCherry revealed that the large UNC-9::GFP plaques reside within or adjacent to the ER (Fig. 5F). We further examined co-localization with HSP-4, another ER-residue protein, and found that the large UNC-9::GFP plaques were resolved into much smaller foci that co-localized with HSP-4 (Fig. 5G), confirming that UNC-9 accumulates in the ER in the absence of *ergi-2* or *ergi-3*.

Notably, the ER-retained UNC-9::GFP aggregates in *ergi-*2 or *ergi-3* mutants appeared in close proximity to degradative organelles, including autophagosomes (labeled by mCherry::LGG-1), late endosomes (labeled by mCherry::RAB-7) and lysosomes (labeled by mCherry::LAAT-1) (Sup. Fig. 2A-D), suggesting that impaired protein degradation may also contribute to aggregate formation.

### ERGI-2 and ERGI-3 facilitate the disassociation of BIP /HSP-4 from IRE-1

Excessive UNC-9 protein accumulates in the ER in *ergi-2* and *ergi-3* mutants, yet the UPR is inhibited. To understand how ERGI-2 and ERGI-3 regulate UPR activation, we sought to identify their interacting proteins. Using CRISPR/Cas9, we generated GFP knock-in strains for *ergi-2* and *ergi-3* and performed anti-GFP immunoprecipitation coupled with mass spectrometry (IP-MS). This revealed a significant overlap in the interactomes of ERGI-2 and ERGI-3 (Fig. 6A and B), supporting their potential function as a complex (Guan et al., 2022). Among the identified partners were components of COPII and COPI vesicles, consistent with their roles in the early secretory pathway (Fig. 6C). Notably, we also detected interactions with proteins involved in ER UPR, redox processes, and lysosomal pathways. Of particular interest was HSP-4/BiP, a central regulator of the ER stress response (Fig. 6C).

**Figure 6:**
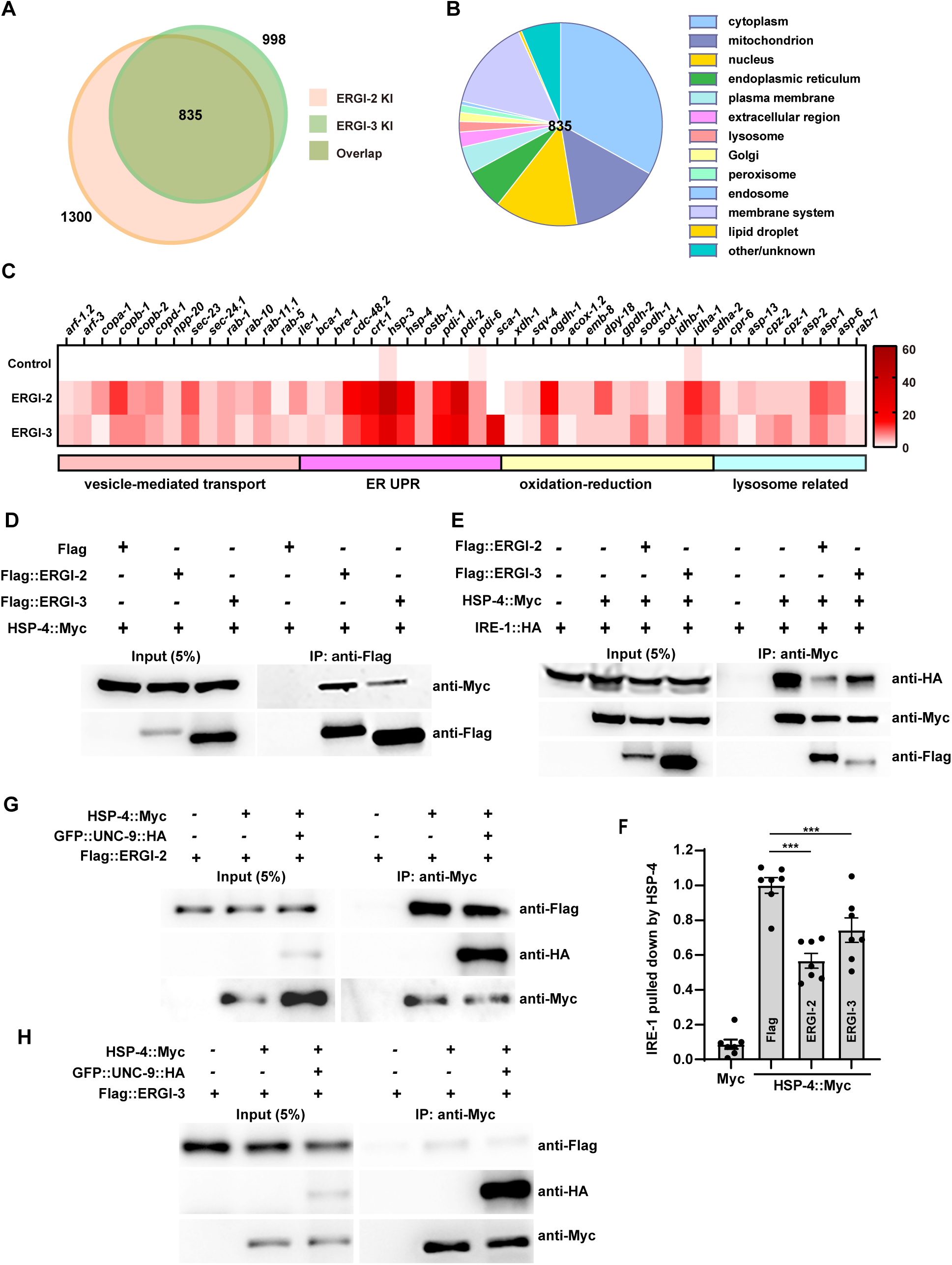
ERGI-2 and ERGI-3 dissociate HSP-4 from IRE-1. (A) Venn diagram of proteins interacting with ERGI-2 and ERGI-3. (B) Subcellular localization analysis of the overlapping ERGI-2/ERGI-3 interactors. (C) Heatmaps of the overlapping ERGI-2/ERGI-3 interactors based on IP-MS results. Wild-type worms expressing GFP were used as a control; GFP::ERGI-2 and GFP::ERGI-3 knock-in (KI) strains were used for the IP-MS analysis. (D) Co-immunoprecipitation showing the association of Flag::ERGI-2 and Flag::ERGI-3 with HSP-4::Myc. (E) Co-immunoprecipitation of IRE-1::HA with HSP-4::Myc in the presence or absence of Flag::ERGI-2 or Flag::ERGI-3. (F) Quantification of the relative amount of IRE-1::HA co-immunoprecipitated with HSP-4::Myc (normalized to HSP-4::Myc and IRE-1::HA input). (G and H) Western blot analysis showing HSP-4::Myc could be co-immunoprecipitated with UNC-9::GFP and Flag::ERGI-2 (G) or Flag::ERGI-3 (H). Blots were quantified using ImageJ. Statistical analysis was performed using one-way ANOVA with Tukey’s multiple comparisons test. Data are presented as mean ± SEM from 7 independent experiments. *P < 0.05; NS, not significant.

Co-immunoprecipitation (Co-IP) assays in HEK293T cells confirmed that both Flag-tagged ERGI-2 and ERGI-3 interact with Myc-tagged HSP-4 (Fig. 6D). Reciprocal Co-IP using anti-Myc antibodies further validated this interaction (Sup. Fig. 3A). ERGI-2 exhibited stronger binding to HSP-4 than ERGI-3, suggesting that ERGI-3 may associate with HSP-4 indirectly via ERGI-2. This interaction is evolutionarily conserved, as human ERGIC2 and ERGIC3 similarly co-precipitated with human BiP (Sup. Fig. 3B). Domain mapping revealed that both the nucleotide-binding domain (NBD) and substrate-binding domain (SBD) of HSP-4 independently interact with ERGI-2, though full-length HSP-4 showed stronger binding (Sup. Fig. 3A), indicating that the intact structure of HSP-4 is required for optimal interaction.

We next investigated how ERGI-2/ERGI-3 binding to BiP/HSP-4 influences UPR activation. IRE1 activation requires BiP dissociation, we then tested whether ERGI-2/ERGI-3 modulate the BiP-IRE1 interaction. Co-IP assays demonstrated that ERGI-2 or ERGI-3 expression reduces HSP-4::Myc binding to IRE-1::HA (Fig. 6E and F). The effect of ERGI-2 was more pronounced, consistent with its stronger affinity for HSP-4. Neither ERGI-2 nor ERGI-3 alone bound IRE-1, suggesting their action is likely HSP-4-dependent (Sup. Fig. 3C).

In addition to UPR activation, ERGI-2 and ERGI-3 bind to UNC-9 and facilitate the ER-to-Golgi transport of UNC-9 protein (Guan et al., 2022). Co-IP experiments further revealed that HSP-4, ERGI-2, and UNC-9 could coexist in a complex (Fig. 6G and H), suggesting ERGI-2/ERGI-3 may cooperate with UNC-9 to displace HSP-4 from IRE-1, thereby activating UPR.

### Restoring UPR alleviates UNC-9 aggregation

We noticed that, in *ergi-2* and *ergi-3* mutants, UPR activation was detectable during early larval development. As shown in Figure 7, P*hsp-4*::mCherry signal highlighting DD/VD cell bodies, although weaker than in wild type, could be observed from L1 through L3 stages. During this period, UNC-9::GFP exhibited a punctate distribution pattern similar to that of wild type (Fig.7A-E). By the L4 stage, however, the P*hsp-4*::mCherry signal in DD/VD neurons was markedly reduced (Fig.7A-E). This decline coincided with a loss of UNC-9::GFP puncta along neurites and the formation of enlarged aggregates in cell bodies (Fig.7A-E). Taken together, these findings suggest that ERGI-2 and ERGI-3 help sustain the UPR, which in turn may prevent the aggregation of UNC-9.

**Figure 7:**
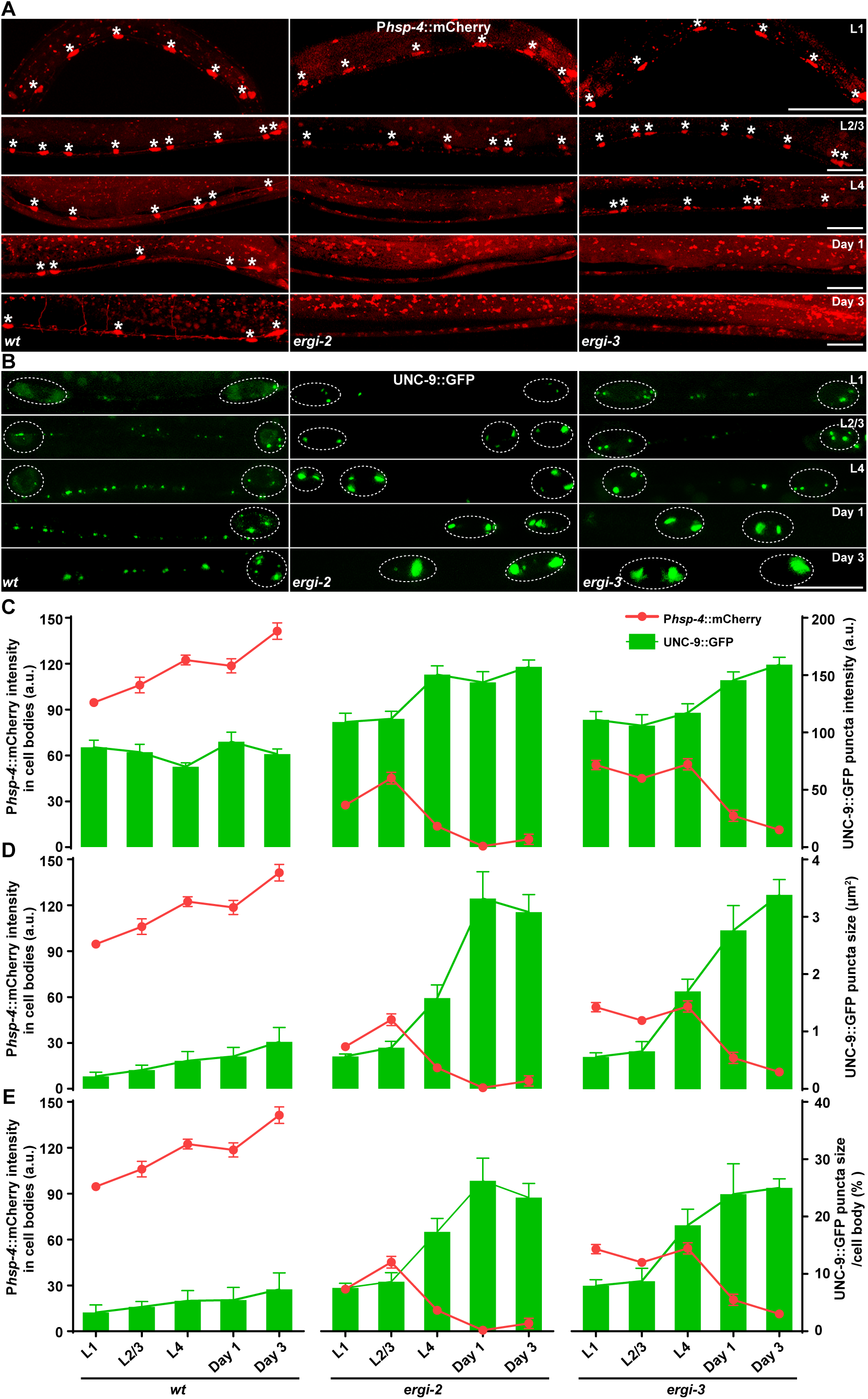
The formation of UNC-9::GFP aggregates is accompanied with the reduced UPR. (A) Expression of Phsp-4::mCherry (red) in DD/VD neurons at different developmental stages (L1, L2/L3, L4 larvae; day 1 and day 3 adults: day1, day3) in wild-type (*wt*), *ergi-2*, and *ergi-3* worms. Asterisks indicate DD/VD cell bodies. Scale bar: 25 µm. (B) UNC-9::GFP distribution in DD/VD neurons at different developmental stages in wild-type (*wt*), *ergi-2*, and *ergi-3* worms. Dashed lines outline the DD/VD cell bodies. Scale bar: 10 µm. (C-E) Quantification of P*hsp-4*::mCherry intensity (red lines) and UNC-9::GFP puncta characteristics in wild-type (*wt*), *ergi-2*, and *ergi-3* worms: (C) average puncta intensity, (D) puncta size, and (E) ratio of puncta area to cell body area.

We therefore asked whether restoring the UPR would alleviate UNC-9 aggregation in *ergi-2* and *ergi-3* mutants. Since the IRE1-XBP1-mediated UPR not only induces chaperon proteins like HSP-4/BiP but also activates autophagy (Guan et al., 2020), we first tested whether overexpressing HSP-4 could resolve the large UNC-9::GFP aggregates. Interestingly, HSP-4 overexpression in DD/VD neurons significantly reduced the size of the abnormally enlarged UNC-9::GFP puncta, and total UNC-9::GFP fluorescence intensity was also decreased (Fig. 8A-E). Consistent with this, double-labeling analyses confirmed that RFP::HSP-4 co-expression reduced UNC-9::GFP aggregate size (Fig. 5G). These results indicate that replenishing chaperone activity is sufficient to mitigate UNC-9 aggregation.

**Figure 8:**
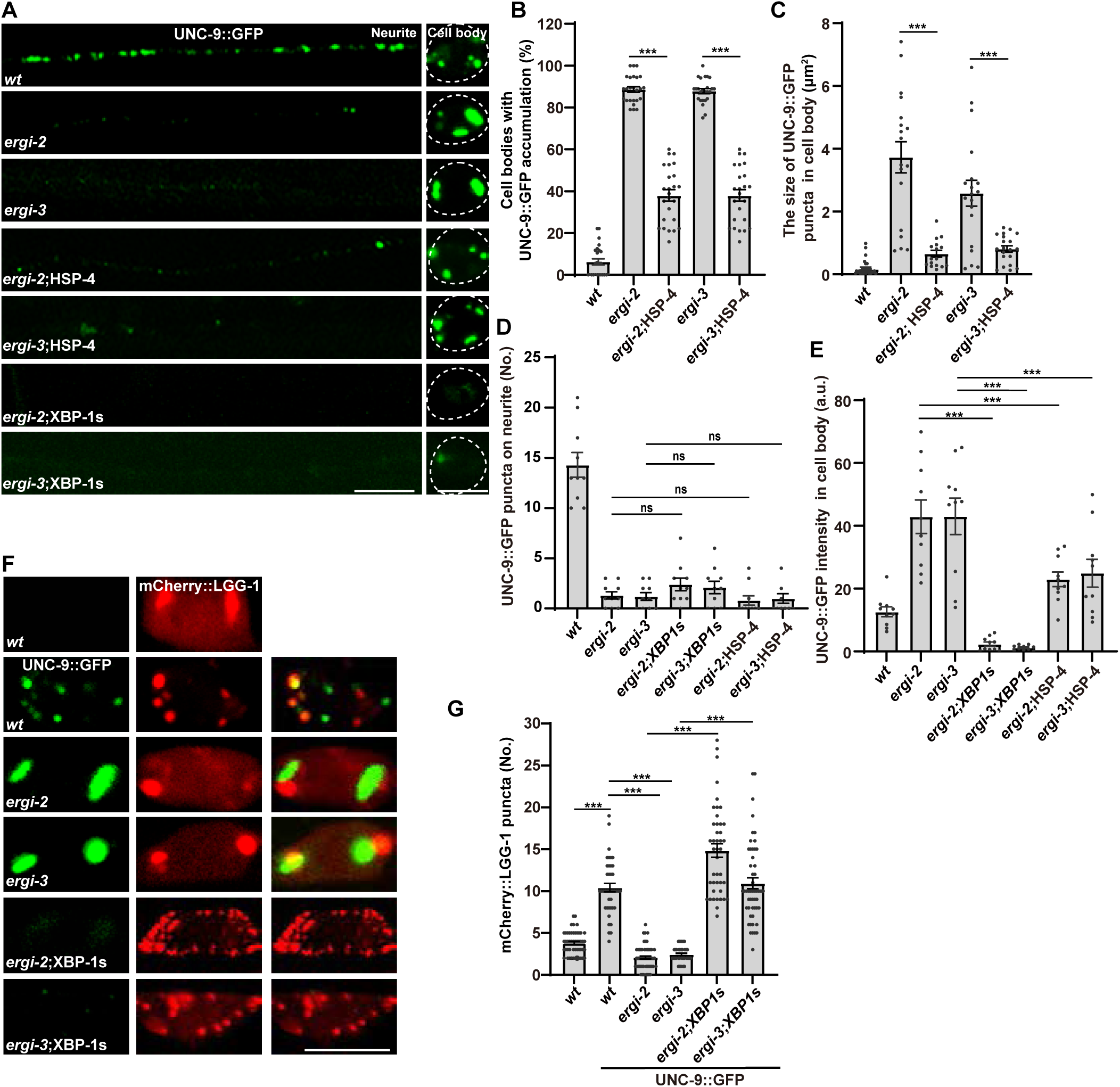
UPR reactivation alleviates UNC-9::GFP aggregates in *ergi-2* and *ergi-3* mutants. (A) UNC-9::GFP (green) aggregates in ergi-2 and ergi-3 mutants are reduced by expression of HSP-4 or XBP-1s. Scale bar: 5 µm. (B-E) Quantification of UNC-9::GFP distribution defects across genotypes. (B) Percentage of cell bodies with abnormal UNC-9::GFP accumulation. (C) Size of UNC-9::GFP puncta in cell bodies. (D) Number of UNC-9::GFP puncta in neurites. (E) UNC-9::GFP intensity in cell bodies. (F) Autophagy activation (mCherry::LGG-1, red) is induced by XBP-1s expression, accompanied by a reduction of UNC-9::GFP (green) aggregates. Scale bar: 5 µm. (G) Quantification of mCherry::LGG-1 puncta number in DD/VD cell body regions. Statistical analysis was performed using one-way ANOVA with Tukey’s multiple comparisons test. Data are presented as mean ± SEM. ***P < 0.001; ns, not significant.

The expression of XBP-1s restores *hsp-4* transcription and enhances autophagy. To determine if this pathway could reduce UNC-9 aggregation, we overexpressed the active form of XBP-1 (XBP-1s) in DD/VD neurons. This intervention effectively suppressed UNC-9::GFP aggregation and nearly eliminated its fluorescent signal in these neurons (Fig. 8A-E). Consistent with this, XBP-1s overexpression also significantly restored autophagy activity in both *ergi-2* and *ergi-3* mutant animals (Fig. 8F and G).

Notably, neither XBP-1s nor HSP-4 overexpression restored UNC-9::GFP signal on DD/VD neurites (Fig. 8A and D), demonstrating that ERGI-2/ERGI-3-mediated transport remains essential for its proper subcellular distribution. Together, these results indicate that, in addition to facilitating ER-to-Golgi transport, ERGI-2 and ERGI-3 proteins promote the activation of IRE1-XBP1-mediated UPR to clear cargo aggregates through chaperone upregulation and autophagy induction.

## Discussion

It is generally considered that the failure of secretory pathway causes abnormal accumulation of cargo proteins in ER, which activates UPR to resolve the protein aggregates in ER (Tang, 2021; Zhang et al., 2021). Here, our findings challenge this paradigm by demonstrating that early secretory components ERGIC2 and ERGIC3 in fact are directly involved in UPR activation in response to cargo overload, revealing a dynamic coupling between protein trafficking and stress signaling.

### The dual role of ERGIC2 and ERGIC3

The dual role of ERGIC2 and ERGIC3 provides a unique opportunity to dissect the relationship between cargo transport and UPR activation. Under normal conditions, cargo proteins are properly folded and efficiently transported out of the ER (Ellgaard and Helenius, 2003; Radanovic and Ernst, 2021), and the UPR remains inactive, as it is not required for cargo localization. ERGIC2 and ERGIC3 bind properly folded cargo proteins and facilitate their export from the ER (Guan et al., 2022). However, when cargo proteins are overloaded, misfolded or excessive cargo saturates ER chaperones (such as BiP), displacing them from stress sensors like IRE1, thereby activating the UPR (Preissler and Ron, 2019). Consequently, the UPR pathway becomes essential for the proper localization of overloaded cargo (Asakura et al., 2015; Shim et al., 2004). Since protein folding is a prerequisite for ER export, the cargo transport function of ERGIC2 and ERGIC3 positions them spatiotemporally close to UPR activation. During cargo overload, ERGIC2 and ERGI3 not only bind to cargo but also interact with BiP/HSP-4, displaying it from IRE1. In this process, overloaded cargo cooperates with ERGIC2/ERGIC3 to dissociate BiP/HSP-4 from IRE1, amplifying UPR activation.

Why are core COPII components not required for cargo-triggered UPR? We suspected that COPII deficiency leads to a global increase of variety of proteins, which is probably sufficient to activate the UPR by dissociating Bip from ER sensors. In contrast, overload of specific cargo may require cargo-specific transport machinery to enhance UPR activation. Given that specific protein accumulations (rather than random protein aggregation) are common in variety of human diseases, particularly neurodegenerative diseases (Hetz and Saxena, 2017), targeting cargo-specific transporters could be a more effective therapeutic strategy. Moreover, in cases of cargo transporter deficiency, activating (rather than inhibiting) the UPR could be an alternative approach to clear toxic aggregates and mitigate disease progression.

In *ergi-2* and *ergi-3* mutants, large protein aggregates are formed adjacent to lysosomes and autophagosomes, suggesting failed clearance. Restoring UPR signaling via XBP-1s overexpression efficiently resolves these aggregates, underscoring the UPR’s critical role in protein quality control. This aligns with studies showing that XBP-1s upregulates lysosomal activity (Imanikia et al., 2019) and that HSP70-family chaperones disassemble neurotoxic aggregates (Kohler and Andréasson, 2023) (Nillegoda et al., 2015). Our work thus identifies ERGIC2 and ERGIC3 as gatekeepers of proteostasis, ensuring that cargo overload triggers adaptive UPR activation rather than toxic accumulation.

### ERGIC2 and ERGIC3 couple cargo transport to UPR activation

Defects in COPII-mediated transport are known to induce ER stress and activate the UPR (Tang, 2021; Zhang et al., 2021). Here, we show that ERGIC2 and ERGIC3 are not merely cargo transporters but are critical regulators of the IRE1-XBP1 pathway. This finding suggests that secretory-machinery components can directly sense cargo burden and modulate UPR signaling. Indeed, several secretory-pathway proteins have been shown to regulate UPR activation by influencing the BiP-IRE1 interaction. For example, the co-chaperone ERdj4 recruits BiP to IRE1, suppressing UPR signaling (Amin-Wetzel et al., 2017), whereases the secretory protein Cab45S binds BiP to inhibit IRE1 activation and apoptosis (Chen et al., 2014). Similarly, the ER-resident factor MANF stabilizes BiP-client complexes and directly interacts with IRE1α to attenuate UPR activity (Kovaleva et al., 2023; Yan et al., 2019). Like these regulators, ERGI-2 and ERGI-3 also modulate UPR activation through BiP-IRE1 interactions, revealing a non-canonical role for COPII-associated proteins in stress response regulation.

Intriguingly, proteomic profiling of purified ERGIC membranes has consistently identified BiP as an abundant resident (Breuza et al., 2004; Kamhi-Nesher et al., 2001), and a previous SEC-23B-centric COPII proteome similarly recovered BiP peptides (Wang et al., 2018; Yehia et al., 2021). This recurrent detection of BiP in early-secretory compartments suggests that ERGIC- and COPII-associated machinery may cooperate to regulate BiP homeostasis, providing independent biochemical evidence for a functional link between early secretory pathway and ER chaperone control.

Notably, ERGI-2 exhibits a stronger influence on UPR activation than ERGI-3, consistent with its higher binding capability for BiP/HSP-4. This functional divergence parallels observations in mammalian cells, where ERGIC2 and ERGIC3 differentially regulate gap junction transport (Guan et al., 2022), and in yeast, where Erv41 and Erv46 play distinct roles in cargo binding (Keiser and Barlowe, 2020). We therefore propose that ERGIC3 primarily mediates cargo recognition, while ERGIC2 additionally serves as a UPR modulator—a dual role that ensures efficient coupling of secretory demand to stress resolution.

### Perspective and future directions

While this study provides mechanistic insight into the role of ERGIC2 and ERGIC3 in coupling cargo transport to UPR activation, several questions remain open for future investigation.

First, gap junction proteins are widely expressed across human tissues. Our findings establish that *ergi-2* and *ergi-3* modulate UPR triggered by neuronal UNC-9 overload. An important next step will be to determine whether their mammalian homologs function similarly in response to the overexpression of gap junction proteins in other tissues or in disease models characterized by gap junction dysregulation.

Second, cargo-receptor pairing is complex and often extends beyond a simple one-to-one relationship. A more systematic mapping of these specific interactions will help clarify how distinct secretory cargos engage the UPR. Additionally, other members of the ERGIC family, such as ERGIC53 and ERGIC32, are known to be essential for the efficient secretion of specific cargos under certain conditions (Appenzeller et al., 1999; Breuza et al., 2004; Itin et al., 1995; Schweizer et al., 1988), even though they are dispensable for bulk flow. It will be valuable to examine whether these related receptors similarly modulate the UPR in response to overload of their respective cargos.

Third, cellular context strongly shapes the response to cargo overload. We observed that UNC-9 overexpression in muscle cells does not induce the UPR, and *ergi-2/ergi-3* are required only for its ER-to-Golgi transport (Guan et al., 2022). In neurons, however, excess UNC-9 activates both UPR and transport pathways dependent on *ergi-2/ergi-3*. Notably, endogenous UNC-9 localization is independent of *ergi-2/ergi-3* and the UPR sensors XBP-1 and IRE-1 (Sup Fig. 4A and B) (Guan et al., 2020). These observations support a model in which the early secretory pathway collaborates with UPR regulators to dynamically manage fluctuations in protein synthesis, folding, and secretion across different cell types and physiological states. Future work should aim to elucidate the precise molecular mechanisms underlying this coordination *in vivo*, including comparative analyses across diverse cells and tissues.

In summary, our work reframes the secretory machinery as an active modulator of UPR signaling, rather than a passive target. By integrating cargo sensing, HSP-4/BiP regulation, and IRE-1 activation, ERGIC-2 and ERGIC-3 provide a mechanistic link between secretory demand and cellular stress resilience. This model offers a promising framework for understanding diseases associated with ER dysfunction, such as neurodegeneration, where defects in cargo handling and UPR dysregulation are prominent pathological features.

## Materials and methods

### Worm strains

Worm strains were maintained on nematode growth medium (NGM) plates at 22°C unless otherwise specified. Culture and genetic manipulation were carried out according to standard procedures (Brenner, 1974). The strains used in this study are listed in Table S1.

### DNA constructs and transgenes

PCR amplifications were performed with KOD-Plus-Neo polymerase (TOYOBO, Cat. #KOD-401) or 2×TransStart FastPfu Fly PCR SuperMix (TransGen, Cat. #AS231-01). Amplicons were cloned into pSM, ΔpSM, pPD95.77, or pPD95.75 vectors via Gibson assembly using the pEASY-Uni Seamless Cloning and Assembly Kit (TransGen Biotech).

Transgenic strains were generated by germline microinjection of the target construct (5-100 ng/μL) with the following co-injection markers: P*myo-2*::RFP (5 ng/μL), P*odr-1*::RFP (50 ng/μL), or P*odr-1*::GFP (50 ng/μL). Extrachromosomal arrays were integrated into the genome by trimethylpsoralen/ultraviolet (TMP/UV) mutagenesis (Gengyo-Ando and Mitani, 2000). Resulting integrants were outcrossed at least three times against the appropriate wild-type background.

### CRISPR/Cas9-mediated gene editing

Knock-in alleles were generated by CRISPR/Cas9-directed homologous recombination. The *xdKi80* (*gfp::ergi-2*) and *xdKi81*(*gfp::ergi-3*) alleles were generated by SunyBiotech and verified by PCR followed by sequencing.

The *xdKi97(wrmScarlet::hsp-4)* allele was engineered using protocols optimized by the Jorgensen laboratory and similarly verified. The targeting sequences and genotyping primers are listed below.

Allele-specific reagents

*xdKi80* (GFP::ERGI-2):

sgRNA: 5′-GCTCGTCGGAAATCCGACAACGG-3′

Forward primer: 5′-TGTAACATTAATATCACCCCAC-3′

Reverse primer: 5′-AAGGGCATTCTGCAATTAGGA-3′

*xdKi81* (GFP::ERGI-3):

sgRNA 1: 5′-CCAATGGACGACTTTCGGGTCAA-3′

sgRNA 2: 5′-GAAAGCCAATGGACGACTTTCGG-3′

Forward primer: 5′-AGAGCAAAGGGCTAACTTAG-3′

Reverse primer: 5′-GCAAGTACAAGTTGACCATAG-3′

*xdKi97* (wrmScarlet::HSP-4):

sgRNAs 1: 5′-ACAGTACGCGTTTGCCACGA-3′

sgRNA 2: 5′-GAAACGCCCAATCAGACGCT-3′

Forward primer: 5′-CAGCCACTCAGCGAACAGTT -3′

Reverse primer: 5′-TCCGGATTGATTGTGAGCTG -3′

### Neuron-specific RNAi

To achieve neuron-specific knockdown, we generated dsRNA from genomic DNA fragments of the target gene coding sequences (0.57–1.2 kb) under the control of the DD/VD neuron-specific promoter P*unc-25* (Esposito et al., 2007; Wu et al., 2025a). The promoter and targeting sequences (50 ng/µL) were amplified and co-injected with the co-injection marker P*myo-2*::RFP (50 ng/µL). The following primers were used to amplify the fragments for *sec-12* and *sec-13* RNAi constructs, which were then recombined into the pPD49.26 plasmid:

*sec-12* front fragment:

Forward primer: CAGGAGGACCCTTGGCTAGCCCAGCTTATTGTCTCAAAAC

Reverse primer: GATATCAATACCATGGTACCCAAATCGAACGCATTTCTGA

*sec-12* rear fragment:

Forward primer: CAGGAGGACCCTTGGCTAGCCAAATCGAACGCATTTCTGAC

Reverse primer: GATATCAATACCATGGTACCCCAGCTTATTGTCTCAAAAC

*sec-13* f front fragment:

Forward primer: CAGGAGGACCCTTGGCTAGCTGGTATGGAATCGAGGATTT TTG

Reverse primer: GATATCAATACCATGGTACCCAAATTGTGATCTGCAAAATT G

*sec-13* rear fragment:

Forward primer: CAGGAGGACCCTTGGCTAGCCAAATTGTGATCTGCAAAAT

Reverse primer: GATATCAATACCATGGTACCTGGTATGGAATCGAGGATTT

### Tunicamycin treatment

Synchronized L4-stage animals were incubated in M9 buffer containing 25 µg/ml Tunicamycin or an equivalent volume of DMSO (1 % v/v final concentration) for 4 hours at 22 °C with gentle agitation. After two washes with M9, worms were transferred to OP50-seeded NGM plates and allowed to recover for 4 hours at 22 °C to permit maximal induction of P*hsp-4*::mCherry.

### Heat-shock treatment

Day 1 adult worms were collected and subjected to heat shock at 30 °C for 5 hours; control animals were maintained at 22 °C. After treatment, worms were immobilized on OP50-seeded NGM plates with 140nM sodium azide, oriented anterior-up under white light, and imaged using a Zeiss Imager A1 microscope controlled by ZEN software (Zeiss).

### Image collection and quantitative analysis

Worms were mounted on 2.5% agar pads in M9 containing 1.4% 1-phenoxy-2-propanol as anesthetic. All fluorescence images were acquired on a Leica SP8 confocal microscope unless otherwise stated. Imaging was performed at the young adult stage unless specified. Confocal stacks were projected to a single image. Comparative images of P*hsp-4:*:mCherry strains were acquired with a 40× objective, while UNC-9::GFP strains were imaged with a 63× objective. Mean fluorescence intensity and co-localization analysis (using Pearson’s correlation) were performed with ImageJ (NIH).

Specific quantitative analyses were performed as follows:

UPR activity in DD/VD neurons: Using the *hsp-4* transcriptional reporter P*hsp-4*::mCherry, we quantified: (1) the number of mCherry-positive somata and commissures, and (2) the average fluorescence intensity in somata and neurites, normalized to wild-type controls.

UPR Induction in Intestine: P*hsp-4*::mCherry fluorescence induced by tunicamycin or heat shock was quantified and normalized to control levels.

Endogenous HSP-4 activation: Using the wrmScarlet::HSP-4 knock-in reporter, we quantified fluorescently labeled neuronal cell bodies.

DD/VD neuron populations: The number of DD/VD neurons was quantified by counting P*unc-25*::mCherry-labeled somata.

*xbp-1* splicing activity: Using the P*unc-25*::XBP-1us::mCherry construct, we counted DD/VD cell bodies highlighted with mCherry signal.

UNC-9::GFP distribution in DD/VD neurons: For strains expressing

P*unc-25*::UNC-9::GFP, we conducted: 1) Line scan analysis: Fluorescence intensity profiles were acquired along the longitudinal axes of cell bodies and neurites using ImageJ and visualized with GraphPad Prism 8.0. 2) Puncta intensity: Mean fluorescence intensities were measured in defined cell body (elliptical ROI, 12 μm²) and neurite (rectangular ROI, 10 μm²) regions. The cell body-to-neurite intensity ratio was then calculated. 3) Puncta number: UNC-9::GFP puncta were counted along neurite segments between the DD5 and VD9 cell bodies. 4) Puncta size: The maximum diameter of individual UNC-9::GFP puncta in cell body regions was measured. 4) Puncta area occupancy: The ratio of the total UNC-9::GFP puncta area to the soma area was calculated to determine cluster coverage.

Endogenous UNC-9 distribution: We used the UNC-9::split-GFP Native and Tissue-specific Fluorescence (NATF) approach to label endogenous UNC-9 (UNC-9-spGFP) (Wu et al., 2025b), combined with P*unc-25*::mCherry to identify DD/VD neurons. The proportion of cell bodies with UNC-9-spGFP signals was quantified.

Autophagy activity: We constructed P*unc-25*::mCherry::LGG-1 and quantified the number of mCherry::LGG-1 foci in DD/VD cell body regions in young adult animals.

### Immunoprecipitation for mass spectrometry analysis

Worm lysates were prepared from the strains *xdKi80* (ERGI-2::GFP) and *xdKi81* (ERGI-3::GFP), using GFP-only expressing worms as a control. Worms were collected and lysed using a Dounce homogenizer in pre-chilled homogenizing buffer (50 mM Tris-Cl, pH 8.0; 150 mM NaCl; 0.5% sodium deoxycholate; 1% Triton X-100; protease inhibitor [Roche]). Lysates were incubated for 15 minutes on ice and centrifuged at 12,000 × g for 15 minutes at 4°C. The supernatants were incubated with a GFP polyclonal antibody (rabbit, Abcam, ab290, 1:1,000 dilution) overnight, followed by incubation with protein A agarose beads (Cat#17-0780-01, GE) for 4 hours at 4°C. The pellets were washed three times with washing buffer (50 mM Tris-Cl, pH 8.0; 150 mM NaCl; 1% NP-40; 1 mM PMSF) and boiled in SDS sample buffer for 5 minutes. The boiled samples were resolved by SDS-PAGE, and protein bands of interest were excised. In-gel digestion was performed using a standard protocol (Shevchenko et al., 2006) with slight modifications. Proteins were reduced with 10 mM DTT, alkylated with 55 mM iodoacetamide, and digested overnight with sequencing-grade trypsin (Sigma) at 37°C. The resulting peptides were analyzed using a TripleTOF 5600 mass spectrometer (AB SCIEX) coupled to an Eksigent nanoLC. Peptide identification and quantification were performed with ProteinPilot 4.2 software (AB SCIEX) using the *C. elegans* proteome sequences (Uniprot) as a database and a mass tolerance of 0.05 Da. The false discovery rate (FDR) was analyzed using the integrated PSPEP software.

Proteins were selected based on significantly elevated or unique Peptide-Spectrum Matches (PSMs) in *xdKi80* and *xdKi81* samples compared to the GFP control. Candidate proteins were subjected to GO enrichment analysis using KOBAS. Functional clusters were visualized via hierarchical clustering and heatmaps (GraphPad Prism 8.0), with color intensity representing relative abundance.

### Immunoprecipitation and western blotting

HEK293T cells were maintained in DMEM supplemented with 10% FBS (Hyclone) at 37°C in 5% CO₂. Transfections were performed using Lipofectamine 2000 (Invitrogen) according to the manufacturer’s instructions. Cells were harvested 24–48 hours post-transfection. For immunoprecipitation, whole-cell extracts were lysed in RIPA buffer (1% Triton X-100, 100 mM Tris-HCl, 50 mM EDTA, 150 mM NaCl, 1% deoxycholate, 0.1% SDS, and protease inhibitor cocktail) for 30 minutes at 4°C. Lysates were centrifuged at 12,000 × g for 15 minutes at 4°C. The supernatants were incubated with anti-FLAG M2 affinity gel or anti-Myc gel beads overnight at 4°C, washed three times with washing buffer, and eluted with SDS-sample buffer. Immunoprecipitated samples or whole-cell lysates were resolved by SDS-PAGE and transferred to nitrocellulose membranes (Pall Life Sciences). Membranes were blocked with 5% non-fat dried milk, and signals were detected using Pierce ECL western blotting substrate (Thermo Fisher Scientific, 34095). All co-immunoprecipitation experiments were repeated at least three times.

### Quantification of protein interactions

To quantitatively assess how ERGI-2 and ERGI-3 modulate the IRE-1::HA and HSP-4::MYC interaction, we used four conditions: (i) Myc-empty vector + IRE-1::HA (negative control), (ii) HSP-4::MYC + IRE-1::HA + empty Flag vector (basal control), (iii) HSP-4::MYC + IRE-1::HA + Flag::ERGI-2, and (iv) HSP-4::MYC + IRE-1::HA + Flag::ERGI-3. Protein band intensities were quantified using ImageJ. HSP-4::MYC immunoprecipitation (IP) efficiency was calculated as the anti-Myc IP/input ratio. IRE-1::HA co-IP efficiency was determined by normalizing anti-HA co-IP signals to the corresponding HSP-4::MYC IP values, adjusting for IP efficiency coefficients, and normalizing against IRE-1 input values. Finally, all interaction ratios were standardized to the Flag-only control group (condition ii) to evaluate the specific effects of ERGI proteins.

## Acknowledgements

We are grateful to Drs. Jianke Gong, Christopher Rongo, Ben Lehner and Yishi Jin for strains and technique support. This work was supported by grants from the National Key R&D Program of China (2024YFA1803401, 2021YFA0805802, and 2024YFA1306100) and the Natural Science Foundation of China (32170790 and 32321004).

## Author contributions

Study concept and design: LG, TZ, ZZ, YW, XH, and MD. Drafting and major editing of original manuscript: MD, LG and TZ. All authors reviewed, revised, and approved the final version of the paper.

## Competing interests

The authors declare no competing interests.

**Correspondence and requests for materials should be addressed to Mei Ding**

## Supplementary Figure Legends

**Sup Figure 1:**
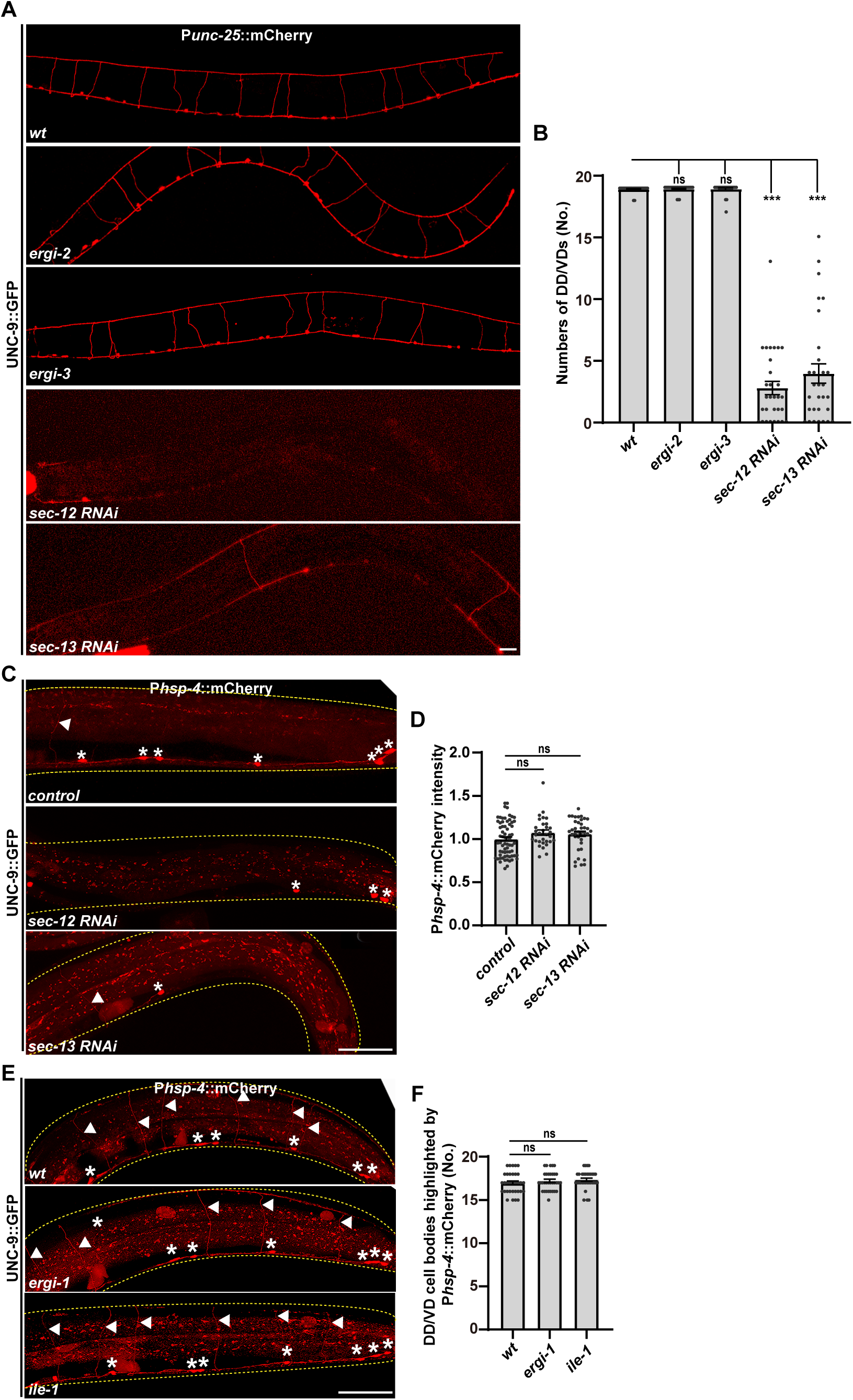
COPII-related components do not affect the UPR. (A) The number of DD/VD neurons (labeled by P*unc-25*::mCherry, red) is decreased in animals treated with *sec-12 RNAi* or *sec-13 RNAi*. (B) Quantification of the number of DD/VD neurons from (A). (C) The UPR (reported by P*hsp-4*::mCherry, red) is unchanged in animals treated with *sec-12 RNAi* or *sec-13 RNAi.* Asterisks indicate DD/VD cell bodies. (D) Quantification of P*hsp-4*::mCherry fluorescence intensity in the remaining DD/VD cell bodies from (C). (E) The UPR (reported by P*hsp-4*::mCherry, red) is unchanged in *ergic-32* and *ile-1* mutant animals. (F) Quantification of P*hsp-4*::mCherry fluorescence intensity in the DD/VD cell bodies from (E). Statistical analysis was performed using one-way ANOVA with Tukey’s multiple comparisons test. Data are presented as mean ± SEM. ***P < 0.001; ns, not significant. Scale bars: 25μm.

**Sup Figure 2:**
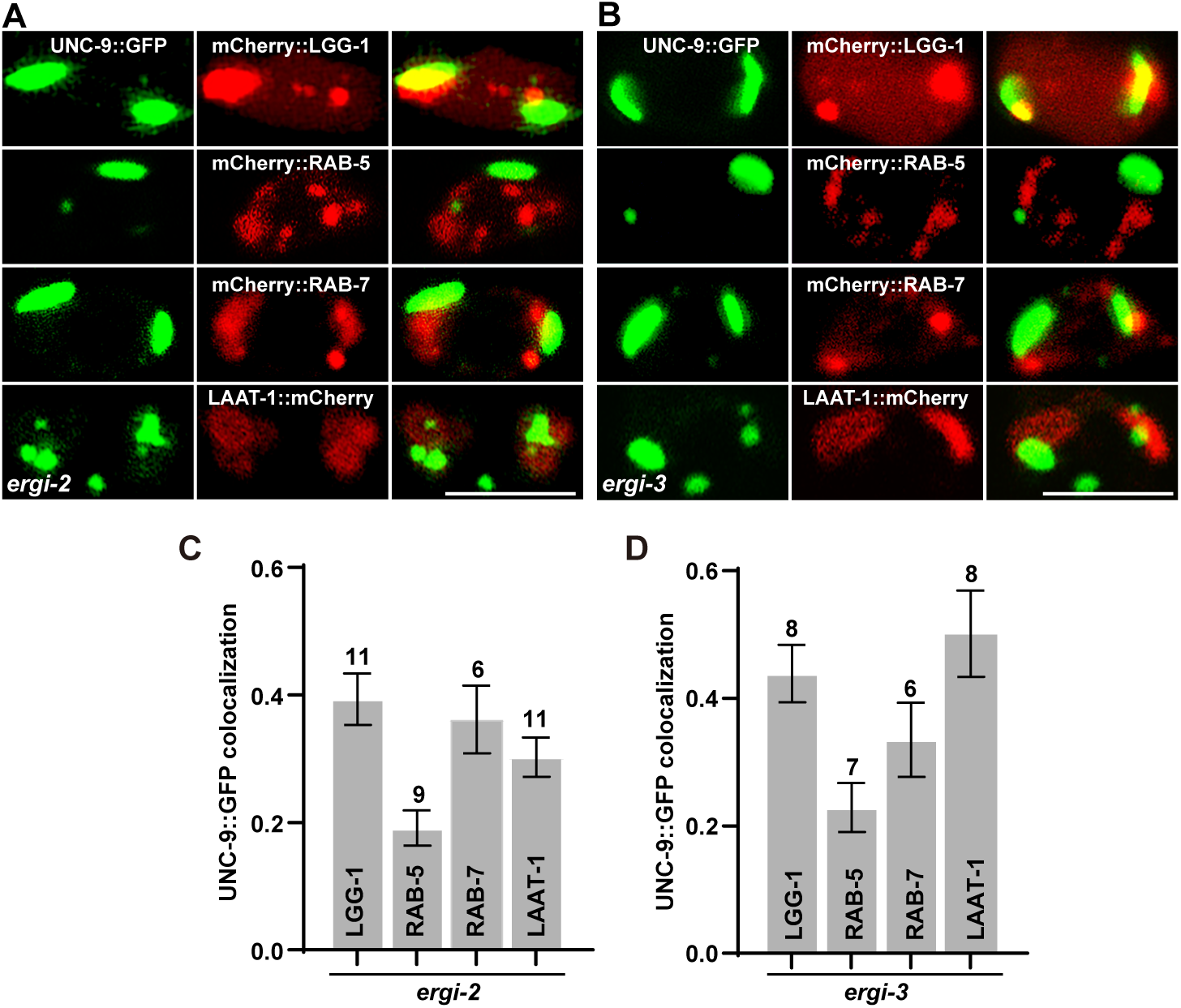
Subcellular localization of UNC-9::GFP in *ergi-2* and *ergi-3* mutants. (A and C) Colocalization analysis between UNC-9::GFP aggregates (green) and autophagosomes (mCherry:LGG-1), early endosomes (mCherry::RAB-5), late endosomes (mCherry::RAB-7) or lysosomes (LAAT-1::mCherry) in DD/VD neurons of *ergi-2* (A) and *ergi-3* (C) mutants. Scale bar: 5μm. (B and D) Quantification of colocalization between UNC-9::GFP and organelle-specific markers in *ergi-2* (B) and *ergi-3* (D) mutants, represented by the Pearson coefficient. Sample numbers are indicated above each column.

**Sup Figure 3:**
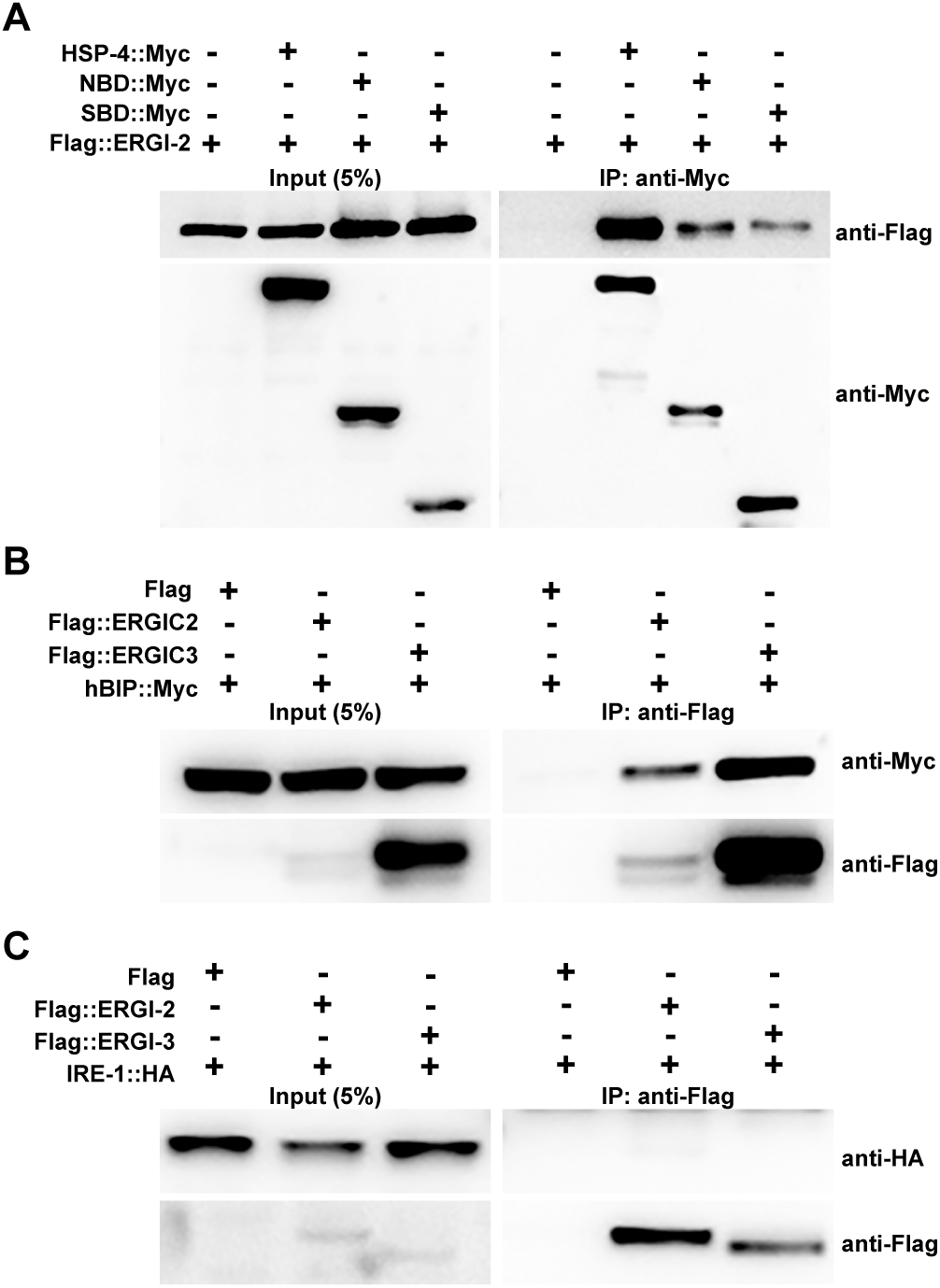
Interactions among HSP-4, ERGI-2, ERGI-3, and UNC-9. (A) HSP-4::Myc co-immunoprecipitates with Flag::ERGI-2. (B) Human ERGIC2 and ERGIC3 co-immunoprecipitate with hBiP. (C) HA::IRE-1 does not co-immunoprecipitate with Flag::ERGI-2 or Flag::ERGI-3.

**Sup Figure 4:**
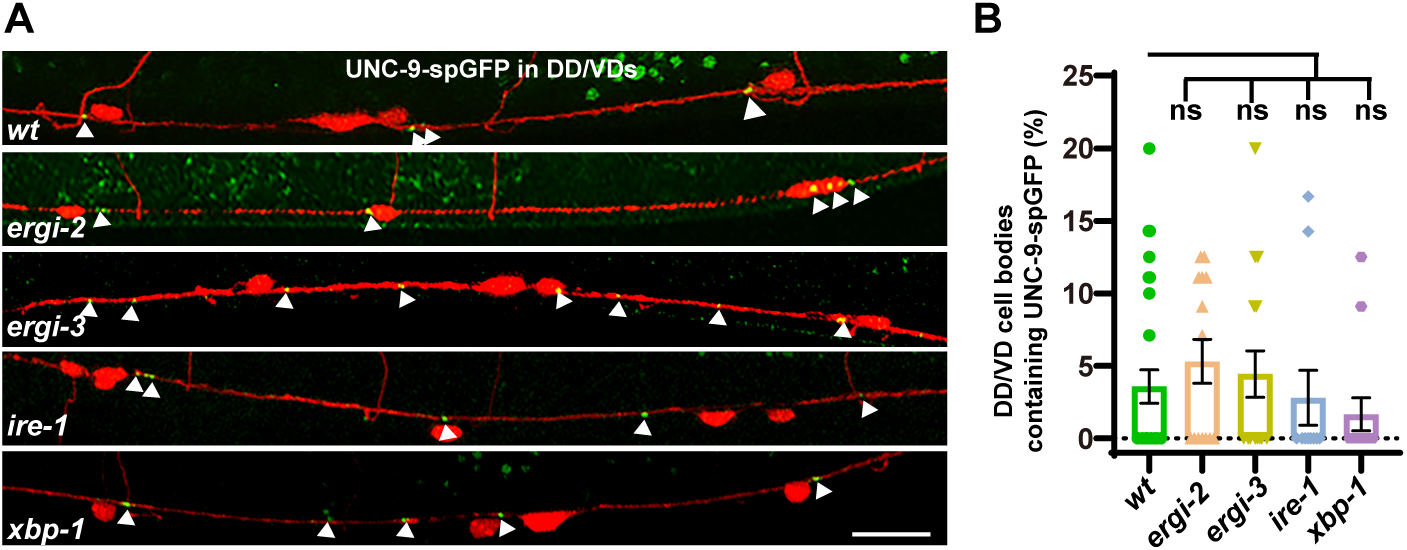
The endogenous UNC-9 distribution is not affected by *ergi-2*, *ergi-3*, *ire-1*, or *xbp-1*. (A) The endogenous UNC-9 (labeled by UNC-9-spGFP, green) in DD/VD neurons (labeled by P*unc-25*::mCherry, red) is not altered in *ergi-2*, *ergi-3*, *ire-1*, or *xbp-1* mutants. (B) Quantification of the proportion of DD/VD cell bodies containing UNC-9-spGFP puncta. Statistical analysis was performed using one-way ANOVA with Tukey’s multiple comparisons test. Data are presented as mean ± SEM. ns, not significant. Scale bars: 25μm.

## References

Adolf, F., Rhiel, M., Hessling, B., Gao, Q., Hellwig, A., Bethune, J., and Wieland, F.T. (2019). Proteomic Profiling of Mammalian COPII and COPI Vesicles. Cell Rep 26, 250–265 e255.

Al-Amin, M., Kawasaki, I., Gong, J., and Shim, Y.-H. (2016). Caffeine Induces the Stress Response and Up-Regulates Heat Shock Proteins in Caenorhabditis elegans. Molecules and Cells 39, 163–168.

Amin-Wetzel, N., Saunders, R.A., Kamphuis, M.J., Rato, C., Preissler, S., Harding, H.P., and Ron, D. (2017). A J-Protein Co-chaperone Recruits BiP to Monomerize IRE1 and Repress the Unfolded Protein Response. Cell 171, 1625–1637 e1613.

Appenzeller, C., Andersson, H., Kappeler, F., and Hauri, H.P. (1999). The lectin ERGIC-53 is a cargo transport receptor for glycoproteins. Nat Cell Biol 1, 330–334.

Araki, K., and Nagata, K. (2011). Protein folding and quality control in the ER. Cold Spring Harb Perspect Biol 3, a007526.

Asakura, T., Ogura, K., and Goshima, Y. (2015). IRE-1/XBP-1 pathway of the unfolded protein response is required for properly localizing neuronal UNC-6/Netrin for axon guidance in C. elegans. Genes Cells 20, 153–159.

Bar-Ziv, R., Frakes, A.E., Higuchi-Sanabria, R., Bolas, T., Frankino, P.A., Gildea, H.K., Metcalf, M.G., and Dillin, A. (2020). Measurements of Physiological Stress Responses in C. Elegans. J Vis Exp.

Barlowe, S.O.a.C. (2002). The Erv41p±Erv46p complex multiple export signals.pdf. The EMBO Journal 21.

Brenner, S. (1974). The genetics of Caenorhabditis elegans. Genetics 77, 71–94.

Breuza, L., Halbeisen, R., Jeno, P., Otte, S., Barlowe, C., Hong, W., and Hauri, H.P. (2004). Proteomics of endoplasmic reticulum-Golgi intermediate compartment (ERGIC) membranes from brefeldin A-treated HepG2 cells identifies ERGIC-32, a new cycling protein that interacts with human Erv46. J Biol Chem 279, 47242–47253.

Burga, A., Casanueva, M.O., and Lehner, B. (2011). Predicting mutation outcome from early stochastic variation in genetic interaction partners. Nature 480, 250-253.

Calfon, M., Zeng, H., Urano, F., Till, J.H., Hubbard, S.R., Harding, H.P., Clark, S.G., and Ron, D. (2002). IRE1 couples endoplasmic reticulum load to secretory capacity by processing the XBP-1 mRNA. Nature 415, 92–96.

Chen, L., Xu, S., Liu, L., Wen, X., Xu, Y., Chen, J., and Teng, J. (2014). Cab45S inhibits the ER stress-induced IRE1-JNK pathway and apoptosis via GRP78/BiP. Cell Death Dis 5, e1219.

Cox, J.S., and Walter, P. (1996). A novel mechanism for regulating activity of a transcription factor that controls the unfolded protein response. Cell 87, 391–404.

Ellgaard, L., and Helenius, A. (2003). Quality control in the endoplasmic reticulum. Nature Reviews Molecular Cell Biology 4, 181–191.

Esposito, G., Di Schiavi, E., Bergamasco, C., and Bazzicalupo, P. (2007). Efficient and cell specific knock-down of gene function in targeted C. elegans neurons. Gene 395, 170–176.

Gengyo-Ando, K., and Mitani, S. (2000). Characterization of mutations induced by ethyl methanesulfonate, UV, and trimethylpsoralen in the nematode Caenorhabditis elegans. Biochem Biophys Res Commun 269, 64–69.

Guan, L., Yang, Y., Liang, J., Miao, Y., Shang, A., Wang, B., Wang, Y., and Ding, M. (2022). ERGIC2 and ERGIC3 regulate the ER-to-Golgi transport of gap junction proteins in metazoans. Traffic 23, 140–157.

Guan, L., Zhan, Z., Yang, Y., Miao, Y., Huang, X., and Ding, M. (2020). Alleviating chronic ER stress by p38-Ire1-Xbp1 pathway and insulin-associated autophagy in C. elegans neurons. PLoS Genet 16, e1008704.

Hauri, H.P., Kappeler, F., Andersson, H., and Appenzeller, C. (2000). ERGIC-53 and traffic in the secretory pathway. J Cell Sci 113 ( Pt 4), 587–596.

Hetz, C., and Saxena, S. (2017). ER stress and the unfolded protein response in neurodegeneration. Nat Rev Neurol 13, 477–491.

Imanikia, S., Ozbey, N.P., Krueger, C., Casanueva, M.O., and Taylor, R.C. (2019). Neuronal XBP-1 Activates Intestinal Lysosomes to Improve Proteostasis in C. elegans. Curr Biol 29, 2322–2338 e2327.

Itin, C., Schindler, R., and Hauri, H.P. (1995). Targeting of protein ERGIC-53 to the ER/ERGIC/cis-Golgi recycling pathway. J Cell Biol 131, 57–67.

Jin, Y., Jorgensen, E., Hartwieg, E., and Horvitz, H.R. (1999). The Caenorhabditis elegans gene unc-25 encodes glutamic acid decarboxylase and is required for synaptic transmission but not synaptic development. J Neurosci 19, 539–548.

Kamhi-Nesher, S., Shenkman, M., Tolchinsky, S., Fromm, S.V., Ehrlich, R., and Lederkremer, G.Z. (2001). A novel quality control compartment derived from the endoplasmic reticulum. Mol Biol Cell 12, 1711–1723.

Keiser, K.J., and Barlowe, C. (2020). Molecular dissection of the Erv41-Erv46 retrograde receptor reveals a conserved cysteine-rich region in Erv46 required for retrieval activity. Mol Biol Cell 31, 209–220.

Kohler, V., and Andréasson, C. (2023). Reversible protein assemblies in the proteostasis network in health and disease. Frontiers in Molecular Biosciences 10.

Kovaleva, V., Yu, L.Y., Ivanova, L., Shpironok, O., Nam, J., Eesmaa, A., Kumpula, E.P., Sakson, S., Toots, U., Ustav, M., et al. (2023). MANF regulates neuronal survival and UPR through its ER-located receptor IRE1alpha. Cell Rep 42, 112066.

Mercado, G., Valdes, P., and Hetz, C. (2013). An ERcentric view of Parkinson’s disease. Trends Mol Med 19, 165–175.

Nillegoda, N.B., Kirstein, J., Szlachcic, A., Berynskyy, M., Stank, A., Stengel, F., Arnsburg, K., Gao, X., Scior, A., Aebersold, R., et al. (2015). Crucial HSP70 co-chaperone complex unlocks metazoan protein disaggregation. Nature 524, 247–251.

Orci, L., Ravazzola, M., Mack, G.J., Barlowe, C., and Otte, S. (2003). Mammalian Erv46 localizes to the endoplasmic reticulum-Golgi intermediate compartment and to cis-Golgi cisternae. Proc Natl Acad Sci U S A 100, 4586–4591.

Plate, L., and Wiseman, R.L. (2017). Regulating Secretory Proteostasis through the Unfolded Protein Response: From Function to Therapy. Trends Cell Biol 27, 722–737.

Preissler, S., and Ron, D. (2019). Early Events in the Endoplasmic Reticulum Unfolded Protein Response. Cold Spring Harb Perspect Biol 11.

Radanovic, T., and Ernst, R. (2021). The Unfolded Protein Response as a Guardian of the Secretory Pathway. Cells 10.

Ron, D., and Walter, P. (2007). Signal integration in the endoplasmic reticulum unfolded protein response. Nat Rev Mol Cell Biol 8, 519–529.

Schindler, R., Itin, C., Zerial, M., Lottspeich, F., and Hauri, H.P. (1993). ERGIC-53, a membrane protein of the ER-Golgi intermediate compartment, carries an ER retention motif. Eur J Cell Biol 61, 1–9.

Schweizer, A., Fransen, J.A., Bächi, T., Ginsel, L., and Hauri, H.P. (1988). Identification, by a monoclonal antibody, of a 53-kD protein associated with a tubulo-vesicular compartment at the cis-side of the Golgi apparatus. J Cell Biol 107, 1643–1653.

Shibuya, A., Margulis, N., Christiano, R., Walther, T.C., and Barlowe, C. (2015). The Erv41-Erv46 complex serves as a retrograde receptor to retrieve escaped ER proteins. J Cell Biol 208, 197–209.

Shim, J., Umemura, T., Nothstein, E., and Rongo, C. (2004). The unfolded protein response regulates glutamate receptor export from the endoplasmic reticulum. Mol Biol Cell 15, 4818–4828.

Sicari, D., Igbaria, A., and Chevet, E. (2019). Control of Protein Homeostasis in the Early Secretory Pathway: Current Status and Challenges. Cells 8.

Stefan Otte, W.J.B., Matthew Heidtman, Jay Liu, Ole N. Jensen, and Charles Barlowe (2001). Erv41p and Erv46p: New Components of COPII Vesicles Involved in Transport between the ER and Golgi Complex. The Journal of Cell Biology 152, 503–517.

Tang, B.L. (2021). Defects in early secretory pathway transport machinery components and neurodevelopmental disorders. Rev Neurosci 32, 851–869.

Walter, P., and Ron, D. (2011). The unfolded protein response: from stress pathway to homeostatic regulation. Science 334, 1081–1086.

Wang, B., Joo, J.H., Mount, R., Teubner, B.J.W., Krenzer, A., Ward, A.L., Ichhaporia, V.P., Adams, E.J., Khoriaty, R., Peters, S.T., et al. (2018). The COPII cargo adapter SEC24C is essential for neuronal homeostasis. J Clin Invest 128, 3319–3332.

Welsh, L.M., Tong, A.H., Boone, C., Jensen, O.N., and Otte, S. (2006). Genetic and molecular interactions of the Erv41p-Erv46p complex involved in transport between the endoplasmic reticulum and Golgi complex. J Cell Sci 119, 4730–4740.

White, J.G., Southgate, E., Thomson, J.N., and Brenner, S. (1986). The structure of the nervous system of the nematode Caenorhabditis elegans. Philosophical transactions of the Royal Society of London Series B, Biological sciences 314, 1–340.

Wu, J., Yarmey, V.R., Yang, O.J., Soderblom, E.J., San-Miguel, A., and Yan, D. (2025a). Heat shock proteins function as signaling molecules to mediate neuron-glia communication in C. elegans during aging. Nat Neurosci 28, 1635–1648.

Wu, Z., Pang, L., and Ding, M. (2025b). CFI-1 functions unilaterally to restrict gap junction formation in C. elegans. Development 152.

Yan, Y., Rato, C., Rohland, L., Preissler, S., and Ron, D. (2019). MANF antagonizes nucleotide exchange by the endoplasmic reticulum chaperone BiP. Nat Commun 10, 541.

Yehia, L., Liu, D., Fu, S., Iyer, P., and Eng, C. (2021). Non-canonical role of wild-type SEC23B in the cellular stress response pathway. Cell Death Dis 12, 304.

Yoshida, H., Matsui, T., Yamamoto, A., Okada, T., and Mori, K. (2001). XBP1 mRNA is induced by ATF6 and spliced by IRE1 in response to ER stress to produce a highly active transcription factor. Cell 107, 881–891.

Zhai, C., and Dong, M.Q. (2025). RME-1-associated recycling endosomes participate in vitellogenin secretion in Caenorhabditis elegans. Life Metab 4, loaf026.

Zhang, Z., Luo, S., Barbosa, G.O., Bai, M., Kornberg, T.B., and Ma, D.K. (2021). The conserved transmembrane protein TMEM-39 coordinates with COPII to promote collagen secretion and regulate ER stress response. PLoS Genet 17, e1009317.

Zhou, L., Zhuo, H., Jin, J., Pu, A., Liu, Q., Song, J., Tong, X., Tang, H., and Dai, F. (2024). Temperature perception by ER UPR promotes preventive innate immunity and longevity. Cell Rep 43, 115071.

